# Functional dissection of human mitotic genes using CRISPR-Cas9 tiling screens

**DOI:** 10.1101/2021.05.20.445000

**Authors:** Jacob A. Herman, Lucas Carter, Sonali Arora, Jun Zhu, Sue Biggins, Patrick J. Paddison

## Abstract

Kinetochores are large protein complexes that assemble at the centromere and bind to mitotic spindle microtubules to ensure accurate chromosome segregation. Like most protein-coding genes, the full multifunctional nature of kinetochore factors remains uncharacterized due to the limited experimental tools for unbiased dissection of human protein sequences. We developed a method that leverages CRISPR-Cas9 induced mutations to identify key functional regions within protein sequences required for cellular outgrowth. Our analysis of 48 human mitotic genes revealed hundreds of regions required for cell proliferation, including known domains and uncharacterized ones. We validated the essential nature for 15 of these regions, including amino acids 387-402 of Mad1, which identified an unknown domain that contributes to Mad1 kinetochore localization and chromosome segregation fidelity. Altogether, we demonstrate that CRISPR-Cas9-based tiling mutagenesis identifies key functional domains in protein-coding genes *de novo*, which elucidates separation of function mutants and allows functional annotation across the human proteome.

## INTRODUCTION

The sequencing of the human genome (Lander et al., 2001) and large-scale genome characterization projects (Consortium, 2012) have provided a reliable list of >20,000 protein coding genes (e.g., UniProt KB database (Breuza et al., 2016)). However, despite this knowledge and intense study of many genes, we lack a comprehensive understanding of how protein activities are compartmentalized into distinct functional domains. Current protein annotation methods mostly rely on homology-based comparative genomic approaches to define domains and infer putative gene functions. For example, the human proteome contains 5494 separate conserved protein family (Pfam) domains each with a putative function (e.g., methyltransferase-like domain) (Mistry et al., 2013); however, >3000 of these domains have unknown function (Bateman et al., 2010) and about half of the proteome is entirely unannotated (El-Gebali et al., 2019; Mistry et al., 2013; Punta et al., 2012). Even within conserved domains experimental validation remains a critical need since homology-based inference is no guarantee for functional similarity. Moreover, large portions of the protein coding genome with critical regulatory functions are inscrutable to homology-based annotation methods because disordered protein regions are only conserved among evolutionarily similar species, yet perform critical cellular functions (i.e., post-translational modifications, recruitment of binding partners, or phase transitions)(Gsponer and Babu, 2009; Ota and Fukuchi, 2017; van der Lee et al., 2014). To this point, within the human genome, long disordered regions are the least likely sequences to be recognized as a Pfam domain (Mistry et al., 2013). Without methods to resolve the sub- functionalization of human proteins independent of homology-based inference we lack the ability to fully characterize these genes.

Current gene manipulation technologies, such as RNAi (Paddison, 2008; Paddison and Hannon, 2002) and CRISPR-Cas9 (Mali et al., 2013; Shalem et al., 2014), although powerful, do not readily resolve the multifunctional nature of proteins coding genes (e.g., catalytic residues, substrate binding interfaces, localization signals, regulatory regions, etc.). Instead, when used in their most common forms, these technologies attenuate *total* gene activity via knockdown, knockout, or transcriptional repression. Thus, they fail to provide adequate insight into a protein’s domain architecture or features. However, one infrequently used CRISPR-Cas9- based approach may be effective at resolving gene domains: tiling sgRNA mutagenesis. This approach was initially employed by two groups attempting to define new design rules for sgRNAs (Munoz et al., 2016; Shi et al., 2015). These studies used pooled sgRNA outgrowth screens where many sgRNAs targeted each protein coding gene to reveal that the sgRNAs causing the most significant changes in outgrowth targeted Pfam domains (Munoz et al., 2016). While these approaches revealed this phenomenon, they did not extend this analysis to identify novel critical protein regions within the coding DNA sequences (CDS). However, this work suggested that sgRNA tiling mutagenesis may reveal the functional landscape of protein coding genes.

Tiling mutagenesis works because in somatic cells, Cas9:sgRNA complexes induce dsDNA breaks that are commonly repaired by error-prone non-homologous end joining (NHEJ), leaving repair scars in the form of small insertion/deletion (indel) mutations (Hartlerode and Scully, 2009; Lieber et al., 2003). Because of the triplet nucleotide nature of protein reading frames, if repair is unbiased, 2/3 of these indels will trigger frameshift mutations, while 1/3 will retain the reading frame (**Fig. 1A**). However, recent deep sequencing of >100 genomic loci in human cells targeted by Cas9 suggest that ∼20% of mutations retain the reading frame (Chakrabarti et al., 2019). Therefore, when using a single sgRNA that targets a *population* of diploid cells, 45-64% of cells should harbor biallelic frameshift mutations, while the remaining cells will carry at least one in-frame, but mutagenized allele (**Fig. 1A**).

**Figure 1.**
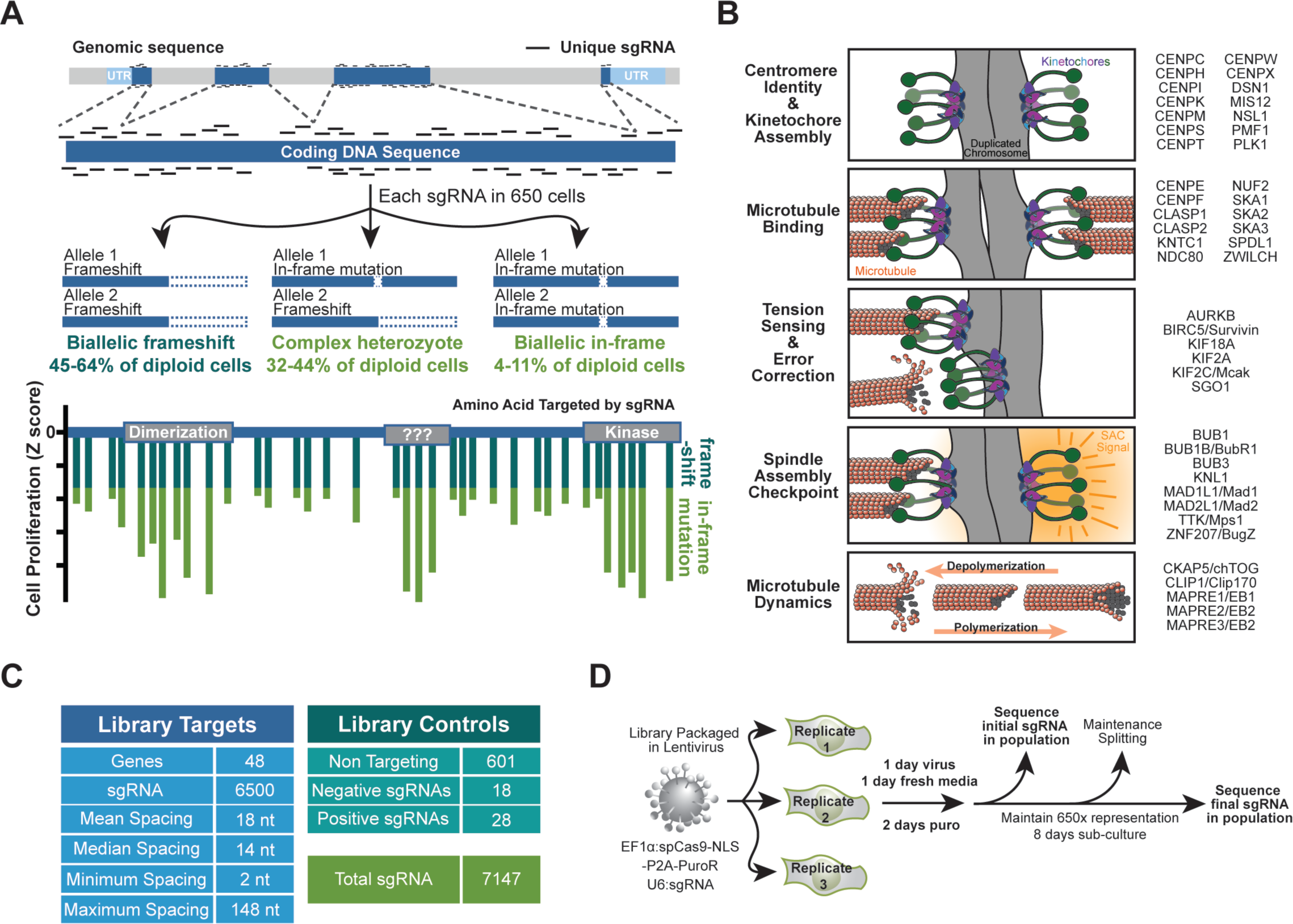
Design and execution of CRISPR/Cas9 tiling screen. (A) Schematic of CRISPR- Cas9 tiling screen and phenotypic readout. The percent of population containing each genotype is calculated assuming unbiased repair (in-frame repairs occur at 33% of edits) or based on deep sequencing (in-frame repairs occur at 20% of edits) (Chakrabarti et al., 2019). (B) Cartoon representation of key molecular activities of kinetochore- and microtubule-mediated processes. Gene/proteins to be screened are listed at right by according to their best characterized function (C) Characteristics of tiling library targeting mitotic factors. (D) Schematic of proliferation-based screening approach using lentiviral particles to insert both spCas9 and sgRNA sequences into genomic DNA. This allows next generation sequencing of sgRNA sequences to serve as an indirect readout of their effect on cell growth.

Thus, sgRNA tiling screens are predicated on the notion that critical protein domains contain amino acid residues that are phenotypically constrained and less mutable than other genic regions. As a result, sgRNAs targeting constrained gene regions will affect the most dramatic phenotypic changes, which will be recognized as “peaks” in the standard next-gen sequence analysis used in outgrowth screens (**Fig. 1A**). A key advantage of this approach is its physiological relevance. Historically, mutagenesis strategies have relied on ectopic overexpression of mutant proteins with unclear physiological relevance, while tiling mutagenesis targets the genomic locus and thus maintains normal cell physiological and protein regulation.

To rigorously test the ability of CRISPR-Cas9 tiling libraries paired with outgrowth screens to elucidate functional protein sequences, we required a set a well-studied, highly multifunctional factors. Such a set of proteins would reveal the ability of proliferation-based tiling libraries to uncover functional motifs that have been previously characterized, independent of homology-based searches. This set would also enable rapid biological characterization of previously unknown functional motifs revealed by tiling. For these reasons we targeted factors that compose or regulate two essential mitotic structures, the microtubule and kinetochore.

Kinetochores are large multi-subunit complexes that are assembled upon centromeres and link chromosomes to the dynamic microtubules of the mitotic spindle. During mitosis, the kinetochore-microtubule attachment physically powers chromosome movements *and* regulates the spindle assembly checkpoint (SAC), which is a biochemical surveillance mechanism that prevents the incorrect segregation of chromosomes (**Fig. 1B**) (Hara and Fukagawa, 2020; London and Biggins, 2014b; Musacchio, 2015). Over three decades of study have revealed much of the underlying chemical and physical properties that enable kinetochore assembly, attachment to microtubules, and SAC surveillance, yet we still do not fully understand the multifunctional nature of these factors.

Here, we selected 48 mitotic genes to target (**Fig. 1B**). These included genes with known rolls in maintaining centromere identity, kinetochore assembly, microtubule binding, tension sensing/error correction, the SAC, and microtubule dynamics (**Fig. 1B**). A comprehensive literature review revealed these proteins contained 167 experimentally defined functional regions and 96 Pfam domains, yet >50% of the coding sequence also fell outside of these areas indicating a chance to reveal novel domains. By performing sgRNA tiling screens in multiple cell lines, we identified hundreds of essential regions among these genes. Approximately 65% of these regions overlap with literature defined functional regions or Pfam domains, while the remaining ∼1/3 of functional regions identified by tiling have not been studied. Consistent with technological limitations associated with interrogating disordered domains, the ‘novel’ functional regions have significant overlap with these rarely interrogate domains. We validated 15 of these functional regions appearing across 6 genes and further characterized the biological role of a previously unknown domain in the SAC protein MAD1L1/Mad1. Thus, the protein regions revealed here are an important resource for future studies of proteins regulating kinetochore and microtubule dynamics, while also representing a key proof of concept for using tiling mutagenesis to dissect multifunctional proteins in mammalian cells.

## RESULTS

### Generation of a CRISPR-Cas9 tiling library targeting mitotic factors

We designed an sgRNA tiling library *in silico* (Methods) that targeted 48 mitotic factors spanning biological functions and genomic contexts (**Fig. 1B**), including two paralogous gene sets, CLASP1/2 and MAPRE1/2/3 (Komarova et al., 2005; Mimori-Kiyosue et al., 2005; Pereira et al., 2006). With genes chosen, we then identified all the *unique* sgRNA targeting the CDS that (with a few exceptions) did not target other coding regions of the genome (**Fig. S1A**). This resulted in a library of 6500 sgRNAs with median spacing of 14 nt between cut sites within the CDS and a maximum spacing of 148 nt due to a lack of the spCas9 protospacer adjacent motif (PAM) (NGG) (**Fig. 1C**).

If the tiling library were to identify functional motifs within a protein, there must be an unbiased distribution of in-frame indels/mutations across a protein’s CDS. Using two different predictors for repair bias after CRISPR-Cas9 editing, we found the library on *average* did not contain any positional bias for sgRNAs predicted to favor frameshifting edits (**Fig. S1B-C**) (Chakrabarti et al., 2019; Shen et al., 2018). However, within a *single gene* some bias could be observed, particularly within short genes that were targeted with only 30-40 sgRNA (**Fig. S1D- E**). Thus, our library should robustly identify essential regions in proteins with more 300 amino acids but may under-report the boundaries or number of functional regions in smaller proteins.

We also included 601 non-targeting control (NTC) sgRNA sequences which cause no editing in the human genome. Thus, NTC sgRNAs reported the rate of unperturbed proliferation to which mitotic-specific sgRNAs were compared (Sanjana et al., 2014). Finally, to monitor screen performance, we also included a small collection of sgRNAs targeting genes that were previously shown to positively (CDKN2A, TP53, etc.) or negatively (POLR2L, HEATR1, etc.) regulate proliferation (O’Connor et al., 2021; Toledo et al., 2015). This final library contained 7147 sgRNAs (**Fig. 1C**) which were synthesized as a pool and inserted into an ‘all-in-one’ single lentiviral expression vector.

To test the effect of each individual sgRNA on proliferation, we infected three independent replicates of cells such that spCas9 and each sgRNA was incorporated into the genome of 650 cells (**Fig. 1D**). The sgRNA sequences were PCR amplified from genomic DNA of populations harvested immediately after infection and after 8 days of outgrowth. Each sgRNA was identified through Illumina-based sequencing to determine how its representation altered over the 8 days of proliferation. The change in normalized sequencing reads for each sgRNA were used to calculate log2(fold change) values and a Z score such that the different cell lines and experimental replicates could be directly compared (**Supplementary Table 1**).

### Tiling proliferation screen is reproducible, and most potent when targeting functional protein regions

To ensure that this approach could be used generally and was not driven by unique cellular or genomic contexts (copy number variations, doubling rates, etc.), we performed the tiling screen in four diverse cell types, which varied in their transformation status, degree of aneuploidy, and tissue of origin. This included common cell lines HeLa (aneuploid) and HCT116 (near diploid), as well as a TERT immortalized retinal pigment epithelial cell line (ARPE^TERT^) (diploid) and laboratory transformed derivative with numerous genetic alterations including an ectopic copy of oncogenic HRAS (ARPE^RAS^) (aneuploid). We found that despite these unique cellular backgrounds, on-target sgRNA (non-targeting controls excluded) affected proliferation similarly in all cell types (**Fig. 2A-B**). The sgRNA affected proliferation in ARPE^TERT^ and ARPE^RAS^ cells extremely similarly (Pearson coefficient 0.96) as expected from their shared lineage, yet sgRNA effects on viability were also strongly correlated (Pearson coefficients > 0.81) in cells from diverse tissue and disease types (**Fig. 2A-B**). These correlations were also observed when only sgRNA with the most potent *decreases* in proliferation were analyzed (bottom quartile) (**Fig. S2A-B**). These results indicated that our techniques were reproducible (data span unique preparations of lentiviral particles, library amplifications, and sequencing runs), but more importantly, that our tiling library had similar *phenotypic* outcomes among diverse cells lines despite the semi-random nature of DNA-damage repair following CRISPR-Cas9 targeting.

**Figure 2.**
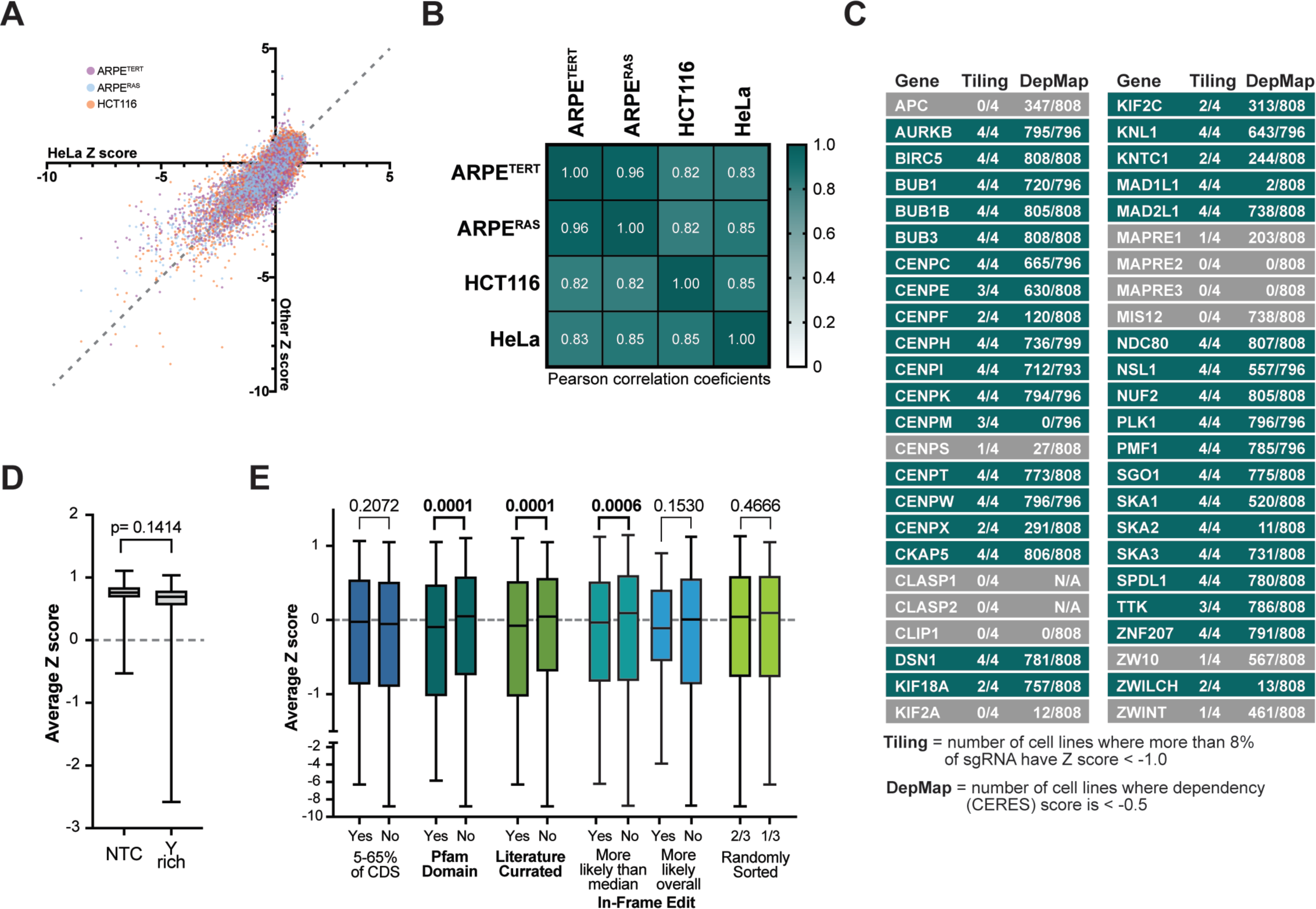
CRISPR/Cas9 tiling screen is technically and biologically reproducible, and sgRNA effects on proliferation are associated with targeting functional protein domains. (A) Each sgRNA’s average Z score from three replicates of ARPE^TERT^, ARPE^RAS^, and HCT116 cells are plotted relative to each sgRNA’s average Z score from three replicates of HeLa cells. These data exclude non-targeting controls, and the dashed line (y = x) is plotted for reference. (B) Correlation matrix and heatmap for all targeting sgRNA in each cell line with Pearson correlation coefficients displayed. (C). Table of 48 genes tiled in the screen, “Tiling” column reports the number of cell lines in which >9% of sgRNA targeting that gene had a Z score < -1.0, indicating that the gene overall was important for cell proliferation. DepMap column reports the number of cell lines screened where a gene is given a ‘dependency score’ < -0.5 at the time of writing, corresponding to a strong negative effect on proliferation. (D) Average Z score for non-targeting controls (NTC) in all four cell lines and targeting sgRNAs that contain the recently reported pyrimidine rich (Y rich) sequence at their 3’ end (Graf et al., 2019). Box plots show median, quartiles, and range of all sgRNAs (E) Targeting sgRNA, minus those containing the Y rich sequence, were binned based on which genomic or protein features they targeted. ‘In- frame edits’ were predicted using Indelphi (Shen et al., 2018) and binned into groups above and below the median likelihood of in-frame edits (22.4%) or above and below the overall likelihood of in-frame edits (50%). Box plots show median, quartiles, and range of the average Z score for each sgRNAs among four cell lines. Mann Whitney tests were used to determine P-values.

The goal of the tiling library was to identify motifs that contribute to the essential activity of mitotic factors, but this first required we determine which of our targets had a negative effect on cell proliferation at the gene-level. We identified all sgRNAs with a Z score less than -1, indicating these sgRNAs had been depleted from the population by at least one standard deviation. Within each gene the percent of sgRNAs meeting this threshold ranged from 0-53% (**Fig. S2C**). Thus, for downstream analysis we excluded the 15% of genes with the least effect on proliferation (fewer than 8% of sgRNAs had Z score less than -1) (**Fig. 2C**). Genes in which more than two cell lines met this threshold are colored teal and reflected trends observed in DepMap studies, which at this time had performed CRISPR screens with 4-6 sgRNA targeting a single gene in more than 750 cell lines (Meyers et al., 2017; Tsherniak et al., 2017).

Our threshold was also consistent with several biological observations, such as targeting CLASP1 or CLASP2 did not affect proliferation because these paralogs function redundantly for most known activities (Mimori-Kiyosue et al., 2005; Pereira et al., 2006). However, there were a few surprising findings, particularly how MAD1L1, MIS12, SKA2, and CENPM behaved differently between our screen and DepMap (**Fig. 2C**). We found that sgRNA targeting the MIS12 gene on *average* failed to meet our gene-level threshold, yet DepMap results showed decreased proliferation in 90% of cell lines after MIS12 targeting (**Fig. 2C**). However, consistent with the DepMap study, the other three proteins in the Mis12 complex (Dsn1, Pmf1, Nsl1) all met our negative proliferation threshold. Looking at the distribution of every sgRNA in our library that targeted MIS12 (**Fig. S2D**), we saw that only four sequences were strongly depleted from the population, and three of those are sequences used in the DepMap library (Sanson et al., 2018). Thus, DepMap identified MIS12 as an essential gene because their library contained primarily the most penetrant sgRNAs, while in the tiling library the signal from these sequences is diluted by the ∼80% of MIS12 targeting sgRNAs that had no effect on proliferation. We hypothesize these sgRNAs failed to cause editing or exhibited repair bias towards the wild-type sequence and thus the gene overall did not meet our threshold. This notion is supported by previous MIS12 inhibition studies that showed lethal chromosome segregation defects in HeLa cells with RNAi knockdown (Goshima et al., 2003).

We observed the reverse behavior in MAD1L1. We found that all four cell lines had negative proliferation outcomes when this spindle assembly checkpoint member was targeted, yet DepMap screening suggests that <1% of cell lines are affected (**Fig. 2C**). We identified four of the six DepMap sgRNA sequences in our data and found that most of those sequences did not affect proliferation, yet with our increased number of sgRNAs we saw many other sequences had a strong negative effect (Sanson et al., 2018)(**Fig. S2C**). We also found that targeting CENPM and SKA2 had negative proliferation outcomes, whereas DepMap data suggest no growth defects (**Fig. 2C**). We realized that this is because DepMap sgRNA sequences for these genes target rarely transcribed exons (**Fig. S2E**). Thus, the high-density data derived for each gene from tiling libraries complement results from other genome-wide approaches and allows interrogation of rare exons and multiple transcripts without confounding the application of the screen at the gene wide level. Altogether, we find that CRISPR tiling is highly reproducible, results in high confidence gene level data, and is rarely limited by biases in CRISPR technology (e.g., inferior performance of MIS12 sgRNAs).

Having identified 36 genes that were required for wild-type levels of proliferation in our data, we set out to determine if any global characteristics drove the performance of the sgRNAs targeting these genes - primarily if sgRNAs targeting functional protein motifs have the most negative effect on proliferation. After synthesizing our library, it was shown that targeting sequences enriched for pyrimidines near the 3’ end in our sgRNA scaffold cause premature polymerase termination (Graf et al., 2019). We identified sgRNAs with these pyrimidine rich (Y rich) sequences within our own data and confirmed those findings independently. Pyrimidine rich sgRNAs on average had no effect on cell proliferation and instead behaved like a non-targeting control (**Fig. 2D**). We excluded these sequences from our global analysis and asked which protein or genomic features were associated with sgRNA activity in our outgrowth screen.

First, we tested conventional wisdom that targeting early exons (within 5-65% of the CDS) resulted in more penetrant loss of activity due to more robust nonsense mediated decay (Doench et al., 2016; Sanson et al., 2018). We found no association between targeting early exons and sgRNA performance (**Fig. 2E**). Instead, our data were consistent with the recent suggestion that sgRNAs targeting functional protein motifs result in the most potent phenotypes because in-frame edits are not tolerated in essential domains (Michlits et al., 2020; Munoz et al., 2016; Shi et al., 2015). We found that sgRNAs targeting Pfam domains or functional regions annotated directly from literature had *on average* the most negative effect on proliferation (**Fig. 2E**). This was promising for our goal of using in-frame mutations to identify functional regions, so we next tested if sites predicted to cause in-frame or frameshifting mutations had the strongest effect on proliferation. In fact, sgRNAs that are predicted to be more likely overall (>50% of cases) or more likely than the median (>22.4% of cases) to create in-frame edits were associated with decreased proliferation (**Fig. 2E**). This strongly suggests that proliferation phenotypes in our screen are *not* driven by frameshift mutations. Together this global analysis is consistent with the observation that in-frame edits caused by repair after CRISPR-Cas9 nuclease activity are common and have the most potent effect on cell proliferation when they occur in an essential region of the CDS.

### Multiple approaches for integrating tiling data within sequence space reveal functional regions

The power of the tiling library is to gain an unbiased understanding of protein function within sequence space. Thus, for each gene we can display the average Z score for all the targeting sgRNAs from the four cell lines (vertical gray bars) along the translated CDS (**Fig. 3A**). In KIF18A, we observe sgRNAs with a strong negative effect on proliferation and sgRNAs that appear largely inactive since they behave similarly to the average of non-targeting controls (**Fig. 3A**). To integrate these data over the CDS we used two previously published approaches, CRISPR-SURF and ProTiler (He et al., 2019; Hsu et al., 2018), that transform results from tiling CRISPR screens into a stepwise function. From this stepwise function, these methods generate ranges of nucleotides or amino acids that are negatively enriched compared to non-targeting controls or a local ‘zero’ (colored regions within each gray bar, **Supplementary Table 2**). As demonstrated by KIF18A, ProTiler and SURF tended to identify many high-resolution regions (10-15 amino acids) that correlate with key functional motifs such as the nucleotide binding pocket in the kinesin motor domain (**Fig. 3A**, pink region near AA 110).

**Figure 3.**
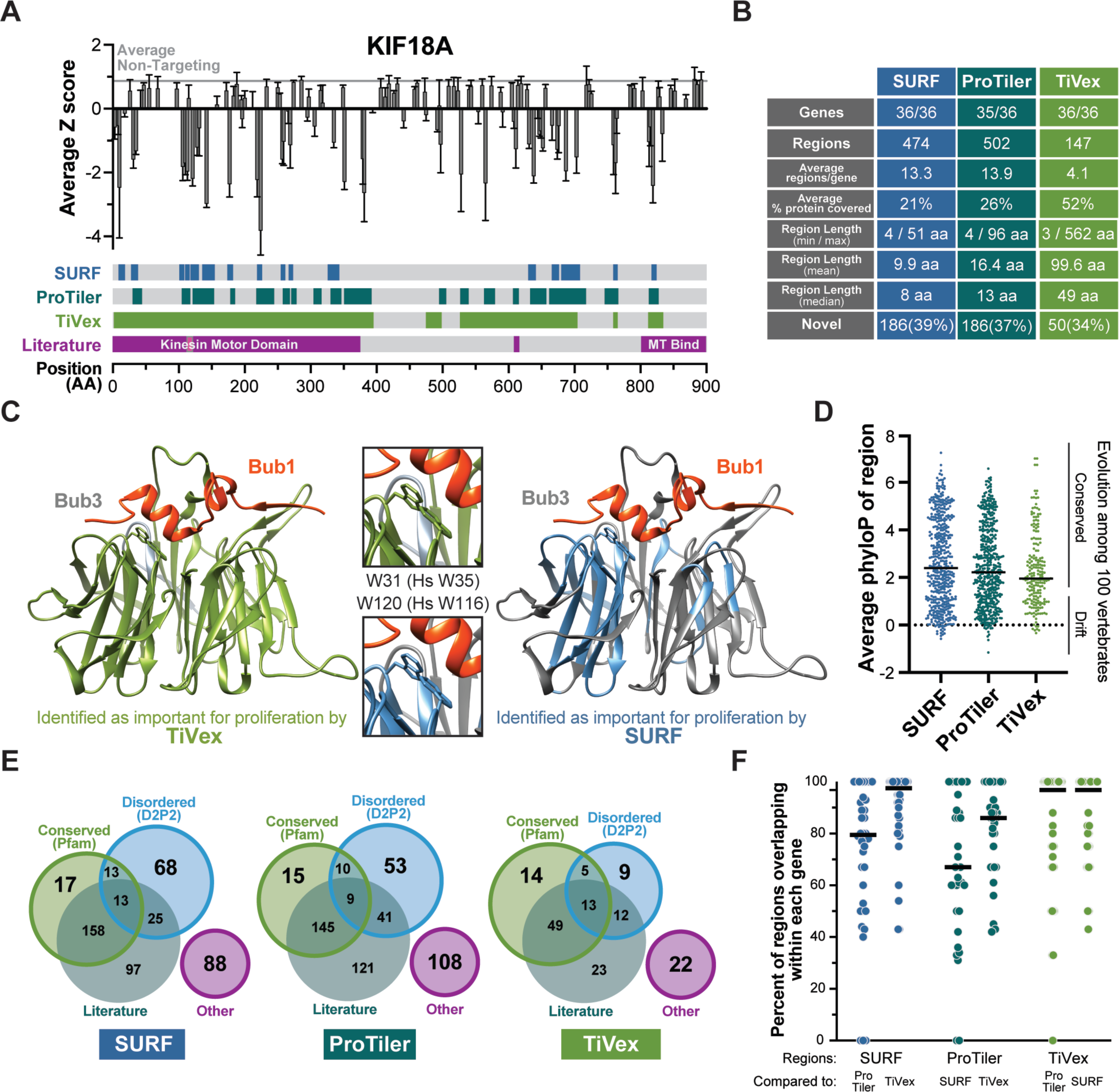
CRISPR/Cas9 tiling data identifies previously known and uncharacterized functional regions including those in evolutionarily divergent and disordered proteins. (A) Median Z score from four cell lines for each sgRNA is mapped to where it targets the CDS of KIF18A (gray vertical bars with 95% confidence interval). Amino acid regions identified by SURF, ProTiler or TiVex as important for cell proliferation are shown as colored in each track corresponding to coordinates at bottom. Kif18A functional regions identified in literature are shown in a similar manner, small pink region within kinesin motor domain highlights the nucleotide binding pocket and the small undescribed region at 610 AA contributes to PP1 binding. Gray horizonal line shows the average value of all non-targeting controls representing normal levels of proliferation. (B) Table describing the characteristics of essential regions identified by analyzing tiling data with SURF, ProTiler, or TiVex in the 36 target genes previously identified (Fig. 2C). (C) Crystal structures of budding yeast Bub3 bound to a peptide of yeast Bub1 (PDB: 2I3S) are colored based on the homologous regions identified by TiVex (green) and SURF (blue) as important for proliferation (Larsen et al., 2007). Two tryptophan residues absolutely required for the protein-protein interaction are shown in zoomed images. (D) PhyloP values (based on the alignment of 100 vertebrate orthologs) for each nucleotide within SURF, ProTiler, or TiVex regions were averaged. Bars represent the median, each dot is a region, and ‘conserved’ corresponds to a value of P < 0.05. (E) Regions identified by SURF, ProTiler, and TiVex, as required for proliferation were grouped by their overlap with conserved Pfam domains (El-Gebali et al., 2019; Mistry et al., 2013), functional regions identified in the literature, or predicted disordered domains (Oates et al., 2013). Regions not meeting fitting these categories are grouped as ‘other’ and represent most uncharacterized regions. (F) Pairwise analysis of overlap between regions identified by each method. Graph displays the number of SURF regions within a single gene (left two columns, blue) that overlaps by at least 1 amino acid with ProTiler or TiVex.

To complement these high-resolution approaches, we paired <u>ti</u>ling data with a con<u>vex</u> fused lasso (TiVex) to generate a more smoothed stepwise function (Parekh and Selesnick, 2015). This allowed TiVex to identify larger windows that overlapped *multiple* SURF or ProTiler regions and often represented discretely folded protein domains such as the KIF18A kinesin domain or CKAP5/chTOG TOG domains (**Supplementary Table 2**) (**Figs. 3A**, **S3A**).

This trend was also evident in an overview of regions identified by each analysis method in our 36 target genes (**Fig. 2C**). SURF and ProTiler identified ∼500 small (10-15 amino acid) regions and TiVex identified ∼150 large (50-100 amino acid) regions (**Fig. 3B**). The resolution of each method can also be demonstrated in three dimensions by mapping SURF or TiVex regions onto the crystal structure of the budding yeast homolog of the target protein Bub3 bound to a fragment of Bub1 (Larsen et al., 2007; Pettersen et al., 2004) (**Fig. 3C**). The entire Bub3 protein folds into a WD40 structure that overall contributes to the interaction with Bub1, thus TiVex identified essentially the entire protein as important for proliferation (**Fig. 3C**, left, green residues). However, SURF primarily identified the two beta sheets that contain a pair of tryptophan residues that are specifically required for Bub1 binding (**Fig. 3C**, right, blue residues) (Larsen et al., 2007; Pettersen et al., 2004). Similarly, mapping TiVex and SURF regions onto the structure of a single domain within chTOG/CKAP5 budding yeast homologue bound to a tubulin dimer showed that TiVex identified the entire fold as important while SURF regions clustered more specifically at the binding interface (**Fig. S3A**).

For some small proteins like Bub3, TiVex identified most of the sequence as important, which is consistent with how the protein functions. We find that on average TiVex identifies ∼55% of the protein sequence within each gene as contributing to proliferation, which is similar to the same analysis with Pfam domains or literature defined functional motifs which cover 50-60% of protein sequence (**Fig. S3B**). The high-resolution nature of SURF and ProTiler are also highlighted by this metric as they identify 20-30% of the overall protein sequence as important for proliferation (**Fig. S3B**). Thus, we propose TiVex may be better suited to identify protein regions that are *sufficient* for a specific activity, while ProTiler and SURF are likely to reveal the regions that are *required* for a specific activity.

TiVex identified protein domains of similar size to Pfam, yet unlike Pfam, TiVex was not restricted to conserved protein sequences. We calculated an average conservation score for each region identified by SURF, ProTiler, or TiVex based on the nucleotide conservation among 100 vertebrates (PhyloP) within the UCSC genome browser (Kent et al., 2002; Pollard et al., 2010) (**Fig. 3D**). Approximately 70% of protein motifs identified in the three analysis methods demonstrated some sequence conservation among vertebrate species (P < 0.05), while the sequence was not constrained in the remaining ∼30% of protein motifs. This suggests that CRISPR tiling screens are applicable to proteins with evolutionarily fluid sequences. When we cross referenced regions identified by SURF, ProTiler, and TiVex with our manually curated list of functional regions identified in literature we found that 34-39% of regions identified by tiling have, to our knowledge, not yet been characterized (**Fig. 3B**). Some of these unstudied motifs overlapped with conserved regions (Pfam domains) but many of them fell in regions predicted to be disordered, or not within either of those categories (**Supplementary Table 3**) (**Fig. 3E**).

Overall, we see strong agreement between all three analysis methods. In pairwise comparisons 100% of the protein regions identified by each method overlap for 7-13 of the genes (**Fig. 3F**) and major discrepancies are primarily focused in 3-4 genes like KNTC1 and CENPF (**Fig. S3C**). These differences likely arise from how each method defines ‘zero’ (relative to NTC or gene averages). Analysis methods are self-consistent but differentially responsive to biological limitations in sgRNA performance.

As a measure of robustness and to test whether sgRNAs with low editing efficiency could obscure important functional motifs, we performed SURF and TiVex analysis on screen data modified to contain low efficiency sgRNA. To this end, we generated new data sets by randomly transforming the Z scores for 10, 20, 30, 40, and 50% of sgRNAs targeting each gene to a value within the range of NTC sgRNAs (**Fig. S4A**). This revealed, in general, that for SURF regions precision and recall scale with the amount of non-functional sgRNAs substitutions, but remain robust at 10% data replacement and that a significant fraction are correctly identified through out (**Fig. S4B,C**). For TiVex domains, substition of non-fucntional sgRNAs were more robust to 20% replacement, likely owing to the larger size of these regions (**Fig. S4D,E**). Overall the analysis suggested that a majority of functional regions are unlikely to be obscured by low efficiency sgRNAs.

Altogether, our analysis further confirmed that sgRNAs most strongly affecting proliferation were correlated with targeting functional protein regions (Munoz et al., 2016; Shi et al., 2015) and the most potent sgRNAs are *not* predicted to favor frameshifting mutations, nor must they target an early exon. Instead, when computational approaches were used to integrate tiling data, we revealed that sgRNA most strongly affecting proliferation were instead concentrated within previously characterized functional protein domains, and a collection of 50- 100 putative new functional regions.

### Biological validations indicated that CRISPR tiling is highly accurate

Because the association of sgRNA depletion with literature defined functional motifs was strong evidence for our approach, we set out to validate a set of uncharacterized functional regions identified by tiling. For this analysis, we selected 15 uncharacterized regions identified among 6 genes (CENPH/Cenp-H, CENPK/Cenp-K, MAD1L1/Mad1, SGO1/Sgo1, SKA3/Ska3, and ZNF207/BuGZ). These included both highly conserved and evolutionarily unconstrained protein regions (**Fig. S5A**). To test these domains, we generated wild type proteins that were resistant to Cas9 editing, and then created mutant proteins that contained small (10-40 amino acid) deletions corresponding to regions identified by SURF, ProTiler, and/or TiVex. These wild type and mutant proteins were N-terminally tagged with a 2xFlag tag and/or EGFP to enable downstream analysis through biochemistry and cell biology, and each one named for the first residue within the small deletion (Ska3^Δ238-253^ shortened to Ska3^238Δ^ or 238Δ). We then used the Flp Recombinase to insert the DNA coding for these exogenous proteins at a unique genomic locus within the parental cell line (Gossen and Bujard, 1992; O’Gorman et al., 1991; Taylor et al., 1998) (**Fig. 4A**). Cell lines encoding the wild type or mutant proteins were then electroporated with Cas9 in complex with 1-2 synthetic targeting or non-targeting sgRNAs (Hoellerbauer et al., 2020a; Hoellerbauer et al., 2020b). Doxycycline was either withheld or added after electroporation to test the effect of endogenous gene knockout and whether expression of the wild type or mutant protein complemented its essential activity.

**Figure 4.**
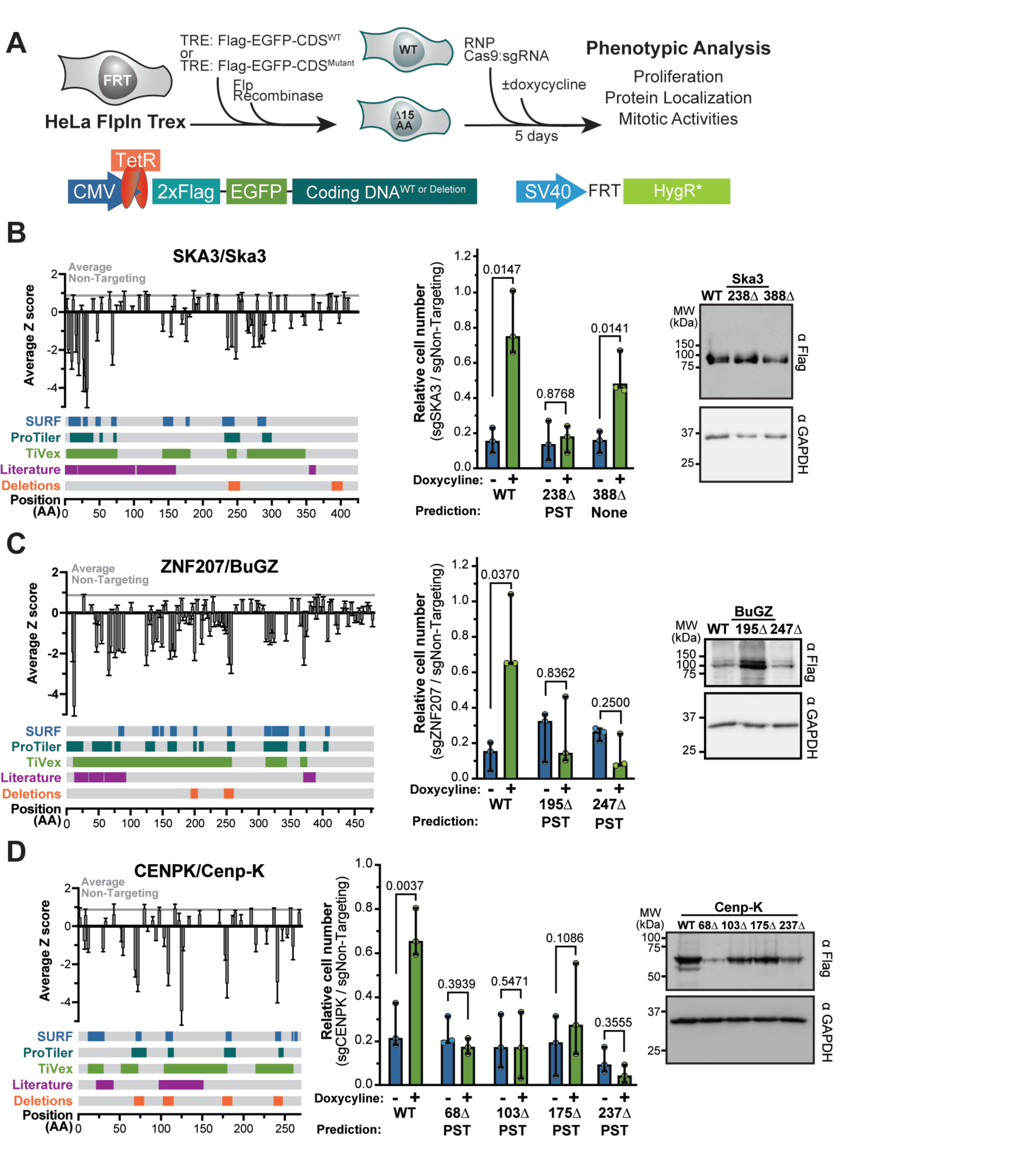
Functional validation and characterization of 11 high-resolution regions within 5 genes identified by tiling. (A) Schematic of generating cell lines in which expression of wild type or mutant proteins is induced by doxycycline and the CRISPR/Cas9-based complementation approach used for functional validation. (B-D) Tiling profile, validation of proliferation phenotype, and assay of protein stability for (B) SKA3/Ska3, (C) ZNF207/BuGZ, and (D) CENPK/Cenp-K. Tiling profiles are the same as Fig. 3 while also showing regions that were deleted. Cell proliferation was assayed as the cell number after knockout of endogenous protein relative to a non-targeting control in the presence (green) or absence (blue) of doxycycline. Cell numbers were normalized to the same cell line electroporated with a non- targeting control. The analysis methods predicting a region to be essential shown as P = ProTiler, S = CRISPR-SURF, and T = TiVex. Each dot is a biological replicate with bars showing median values and 95% confidence intervals. Paired t tests were used to determine P-values. Steady state protein levels of wild type and mutant proteins were assayed by immunoblot in the presence of endogenous protein, using GAPDH as a loading control and exposed for a shorter interval. (More examples in Fig. S4).

We tested regions within Ska3, BuGZ, and Cenp-K that were identified by all three computational methods, and one additional region in Ska3 that was not identified in the screen as a control (**Fig. 4B-D**). In all cases, wild type proteins provided a significant rescue for cell proliferation following endogenous protein knockout, as did the control deletion in Ska3 (**Fig. 4B- D**). We observed the same behavior in Cenp-H and Sgo1 deletion mutants that were predicted by all three methods, but also found that a region identified solely by ProTiler was a false positive and was not required for proliferation in our validation study (**Fig. S5B-C**). Altogether, using this complementation approach we verified that 10/11 ProTiler regions, and 10/10 regions overlapping with SURF and TiVex windows were required for cell proliferation. This comprehensive analysis suggests that CRISPR-Cas9 tiling libraries are a reliable means to identify previously uncharacterized protein regions.

### Tiling MAD1L1/Mad1 reveals a motif that contributes to its kinetochore localization

Our initial validation focused on some of the most robust regions predicted by all three analysis methods, so next we validated a case where analysis methods showed less agreement, the MAD1L1 gene. Consistent with previous literature, SURF, ProTiler, and TiVex all agreed that the C-terminus of the protein is particularly important for its essential activity (**Fig. 5A**). This region contributes to Mad1 forming a homodimer and then binding to kinetochore factors like Bub1 and Cdc20 (Allan et al., 2020; Brady and Hardwick, 2000; Fischer et al., 2021; Kim et al., 2012; Lara-Gonzalez et al., 2021; London and Biggins, 2014a; Piano et al., 2021). However, in the 600 amino acids upstream of that region we saw very little agreement between SURF, ProTiler, and TiVex (**Fig. 5A**). Thus, we generated four deletion mutants outside the well characterized C-terminus that were identified by only SURF or SURF and TiVex (**Fig. 5A**). Using the same approach as before (**Fig. 4A**), we tested the ability of mutant proteins to complement MAD1L1 knockout. Consistent with previous observations (Allan et al., 2020; Rodriguez-Bravo et al., 2014), the Mad1 protein was long lived and complementation assays could only be performed 10 days after Cas9:sgRNA transfection resulting in greater variability for this assay. Nevertheless, Mad1^WT^ and Mad1^170Δ^ partially rescued the proliferation defect, while mutants Mad1^25Δ^ and Mad1^272Δ^ that were identified by SURF and TiVex did not, recapitulating screen results (**Fig. 5B**). Mad1^387Δ^ which was identified only by SURF rescued viability but with much more variability (**Fig. 5B**). The 10 days required to deplete Mad1 protein led to high variability in proliferation assays that would confound more nuanced mitotic phenotypes, so we further interrogated the biological functions of these essential regions in the presence of endogenous Mad1 protein, as has been done by others (Kim et al., 2012). We validated that none of the mutations compromised protein stability (**Fig. 5C**) and then determined if mutant proteins were able to perform an essential Mad1 activity: maintaining the spindle assembly checkpoint. We induced expression of each Mad1 protein overnight and then treated cells with the microtubule destabilizing drug nocodazole for 20 hours, which should trigger a robust SAC arrest. However, we found that fewer cells expressing Mad1^387Δ^ arrested in mitosis following this treatment, indicating this region of Mad1 contributes to SAC signaling (**Fig. 5D**).

**Figure 5.**
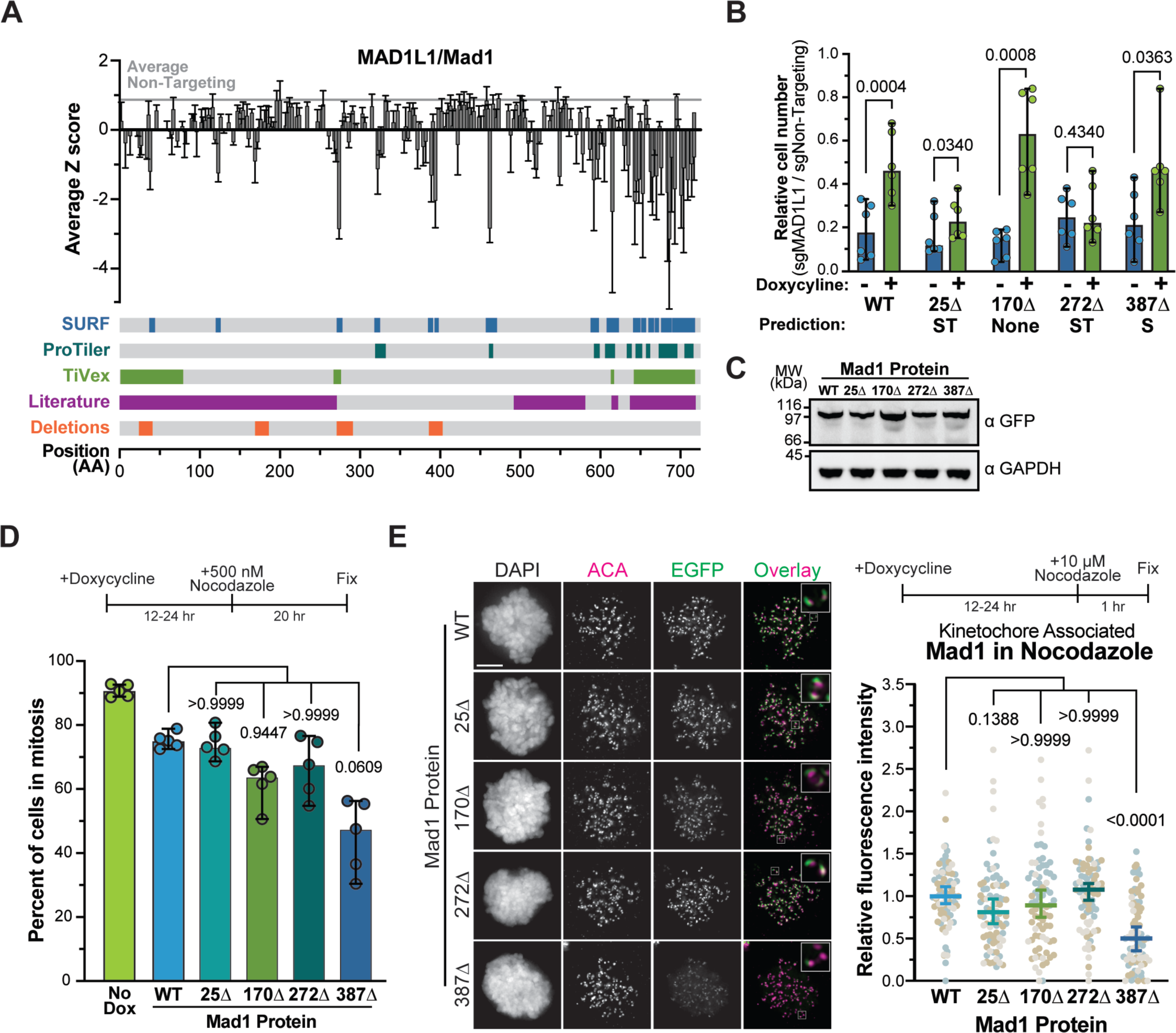
Tiling MAD1L1/Mad1 reveals a region contributing to prolonged activation of the SAC and kinetochore localization. (A) Tiling profile for MAD1L1/Mad1, displayed the same as Fig. 4. (B) Normalized cell number after knockout of endogenous protein in the presence (green) or absence (blue) of doxycycline in each cell line. Dots represent 3 biological replicates performed in duplicate with paired t tests used to determine P-values. (C) Steady state protein levels of wild type and mutant proteins were assayed by immunoblot in the presence of endogenous protein, using GAPDH as a loading control and exposed for a shorter interval. (D) The ability wild type or mutant proteins to maintain a prolonged SAC arrest in the presence of endogenous Mad1 was assayed by treating cells with nocodazole for 20 hours and determining the percent of cells in mitosis based on chromatin morphology. Data in ‘No Dox’ are determined from all five cell lines not exposed to doxycycline. Each dot represents a biological replicate and Dunn’s multiple comparisons test was used to determine P-values. (E) Kinetochore association of EGFP-Mad1 wild type and mutant fusion proteins was determined by the EGFP signal proximal to anti-centromere antibody (ACA) in the presence of endogenous Mad1 and nocodazole. Representative images on left with quantifications on right. Each dot represents the average kinetochore signal from a single cell, cells from three biological replicates are colored differently. Dunn’s multiple comparisons test was used to determine P-values. Scale bars are 1 µm, all averages and error bars in figure are median values and 95% confidence intervals.

Robust SAC signaling requires that Mad1 localize to the kinetochore where it serves as a scaffold to assemble the biochemical inhibitor of mitotic progression (Brady and Hardwick, 2000; De Antoni et al., 2005; Lara-Gonzalez et al., 2021; Piano et al., 2021). Thus, we assayed the ability of mutant proteins to localize to kinetochores in cells either normally transiting mitosis or those experiencing a robust SAC signal due to nocodazole treatment. We found that only the Mad1^387Δ^ protein exhibited kinetochore localization defects, and this occurred specifically when cells were treated with nocodazole (**Figs. 5E and S6**). In these cells, Mad1^387Δ^ kinetochore levels were reduced, yet a significant amount of protein still localized, indicating at least one kinetochore recruitment mechanism remained functional in this mutant.

### Mad1^387Δ^ and Mad1^R617A^ contribute to kinetochore recruitment independently

Recent evidence suggests that Mad1 is initially recruited to kinetochores by the protein Bub1, but when kinetochores remain unattached to microtubules for long periods (such as in nocodazole) the RZZ complex (Rod, Zw10, Zwilch) recruits a separate population of Mad1 to kinetochores (Kim et al., 2012; Rodriguez-Rodriguez et al., 2018; Silio et al., 2015; Zhang et al., 2015). Thus, we hypothesized that Mad1^387Δ^ exhibited kinetochore localization defects specifically in nocodazole because this region contributes to an interaction with RZZ. Our hypothesis would also explain why this region gave mixed results in the proliferation retest: in normally cycling cells the Bub1 pathway is sufficient for SAC activity, while RZZ-recruitment is only required when chromosome alignment defects occur.

Thus, to distinguish between the Bub1 or RZZ recruitment pathways we also inhibited the well characterized Bub1 binding ‘RLK motif’ in Mad1. Mutating Arginine 617 to Alanine in Mad1 (Mad1^R617A^) prevents its biochemical association with Bub1 and reduces kinetochore localization in cells (Brady and Hardwick, 2000; Fischer et al., 2021; Kim et al., 2012; Zhang et al., 2015). We therefore generated the Mad1^R617A^ mutant alone or in combination with Mad1^387Δ^ to determine if mutating both regions entirely prevented kinetochore recruitment (**Fig. 6A**). We overexpressed these mutant proteins and found that fewer cells were able to maintain a SAC arrest in mitosis when expressing Mad1^387Δ^ or Mad1^R617A^ versus Mad1^WT^ (**Fig. 6B**). When the mutations were combined, we observed a slight additive effect, but we suspect this was limited by the presence of endogenous Mad1 protein (**Fig. 6B**). Consistent with the loss in SAC activity observed when either Mad1^387Δ^ or Mad1^R617A^ are over-expressed, we found that both mutations compromised Mad1 association with kinetochore by ∼50% after one hour of nocodazole treatment (**Fig. 6C**). When the mutations were combined, the protein virtually failed to localize to kinetochores. Consistent with previous results (Kim et al., 2012), this suggests that neither Mad1^387Δ^ nor Mad1^R617A^ dimerize with endogenous protein or that such dimers fail to bind kinetochores. More importantly, this indicates that Mad1 residues 387-402 contribute to its kinetochore localization, in a manner that is likely independent of the Bub1 interaction. Thus, this region may mediate or stabilize an interaction with RZZ or another fibrous corona member.

**Figure 6.**
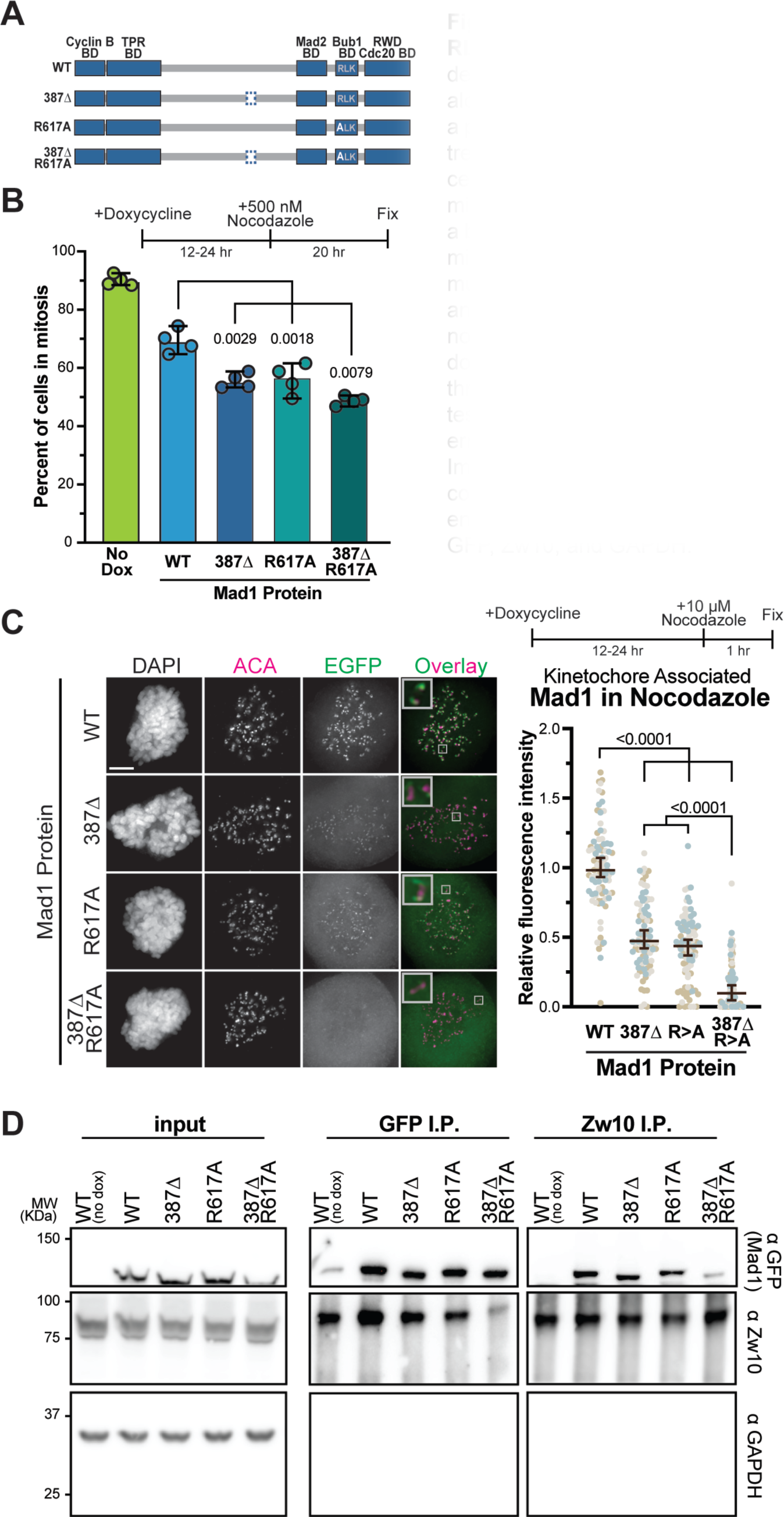
Mad1^387Δ^ contributes to kinetochore localization independent of RLK motif that mediates Bub1 interaction. (A) Schematic of Mad1^387Δ^ deletion mutant (removal of amino acids 387-402) and Mad1^R617A^ point mutant alone or in combination. (B) The ability wild type or mutant proteins to maintain a prolonged SAC arrest in the presence of endogenous Mad1 was assayed by treating cells with nocodazole for 20 hours and determining the percent of cells in mitosis based on chromatin morphology. Data in ‘No Dox’ are determined from all four cell lines not exposed to doxycycline. Each dot represents a biological replicate and Dunn’s multiple comparisons test was used to determine P-values. (C) Kinetochore association of EGFP-Mad1 wild type and mutant fusion proteins was determined by the EGFP signal proximal to anti- centromere antibody (ACA) in the presence of endogenous Mad1 and nocodazole. Representative images on left with quantifications on right. Each dot represents the average kinetochore signal from a single cell, cells from three biological replicates are colored differently. Dunn’s multiple comparisons test was used to determine P-values. Scale bars are 1 µm, all averages and error bars in figure are median values and 95% confidence intervals. (D) Immunoblots showing co-immunopurification of Mad1 proteins and RZZ complex member, Zw10. The same cell lysates (left) were used to purify exogenous EGFP-Mad1 (middle) or endogenous Zw10 (right) and then probe for GFP, Zw10, and GAPDH.

To test this, we asked if endogenous Zw10 co-purified with EGFP-tagged Mad1 proteins from a population of cells arrested in mitosis by treating them with MG132 and nocodazole for four hours. In these biochemical assays, co-purification of Zw10 with Mad1^387Δ^ and Mad1^R617A^ was inconclusive, while the double mutant showed a robust loss of co-purifying Zw10 (**Fig. 6D**). Consistent with this notion, when the immunopurification experiment was reversed to isolate endogenous Zw10, the double mutant largely failed to co-purify (**Fig. 6D**). While we cannot conclude whether Mad1 387-402 and the Bub1 binding RLK motif function entirely independently, it is evident that Mad1 387-402 facilitates an interaction with the RZZ complex. In addition, this result highlights the utility of our sgRNA tiling approach to identify gene domain activities and reveal separation of function mutants.

## DISCUSSION

We have demonstrated that CRISPR-Cas9 tiling mutagenesis of endogenous protein coding sequences in the human genome can be used to functionally validate and identify critical protein regions, including conserved and divergent protein sequences. Our approach takes advantage of the naturally occurring mutagenic properties of error-prone NHEJ in human cell lines after a dsDNA break is introduced by Cas9 activity. From 48 kinetochore-associated genes, we found that 36 produced reliable results for our phenotypic readout (i.e., cell growth) (**Fig. 2C**), which could be represented as unique sgRNA tiling profiles for each (**Fig. 3-4**). The negatively enriched sgRNA regions from these screens, we argue, represent differentially constrained regions of the encoded protein, which are less mutable than other regions due to the presence of an essential gene activity.

We can imagine three explanations for why these regions are sensitive to mutations: that they harbor a critical protein-protein interaction or catalytic activity; that they direct protein folding or stability (Labbadia and Morimoto, 2015); or that they contain non-coding DNA elements (e.g., transcriptional or splicing enhancers) embedded in the protein coding sequence. Our correlations between sgRNA performance and literature defined functional domains (**Figs. 2-3**) and our complementation studies in cells (**Figs. 4, S5**) are consistent with the first two explanations. Our results strongly suggest that protein folding/stability explains a minority of functional motifs identified through tiling as ∼70% of the regions were previously documented in literature to perform enzymatic or binding functions (**Fig 3B,E**). Moreover, in our analysis of uncharacterized motifs we observed protein stability phenotypes in only 3 of 15 mutant proteins that we tested. Thus, it is likely tiling mutagenesis of other gene sets will primarily reveal unique protein-protein interactions or enzymatic behaviors.

In the process of validating CRISPR-Cas9 tiling as a discovery tool, we also generated a powerful resource for the study of kinetochore genes. This includes a rich dataset of experimentally validated sgRNA sequences, but more importantly, while kinetochore factors were the subject of robust and groundbreaking study for nearly four decades, our tiling screen paired with three complementary domain calling methods (i.e., SURF, ProTiler, and TiVex) identified 50-186 essential regions in 36 kinetochore proteins that have not yet been studied (**Fig. 3B, E**). Previous efforts to dissect human kinetochore factors relied on structure or sequence homology to guide truncations or mutations, but our functional screening was not limited in this way (**Fig. 3D-E**). Revealing important regions that would otherwise take years of lab work to identify expedites our collective molecular understanding of kinetochore biology.

The power of CRISPR-Cas9 tiling was also demonstrated by our analysis of a previously unstudied region in MAD1L1/Mad1. Mad1 localization to the kinetochore is dependent on interactions with Bub1 and the RZZ complex (Kim et al., 2012; Rodriguez-Rodriguez et al., 2018; Zhang et al., 2015). While the interaction with Bub1 has been deeply characterized (Brady and Hardwick, 2000; Kim et al., 2012; Lara-Gonzalez et al., 2021; Piano et al., 2021; Silio et al., 2015), through tiling mutagenesis, we identified a 16 amino acid sequence in the middle of the Mad1 protein (aa 387-402) that contributes to its SAC activity, kinetochore localization, and interaction with the RZZ complex. Our findings are consistent with studies that indicate two populations of Mad1 exist at the kinetochore and they rely on distinct regulatory mechanisms (Kim et al., 2012; Rodriguez-Rodriguez et al., 2018; Zhang et al., 2019; Zhang et al., 2015). Moreover, three-dimensional mapping of kinetochore organization within cells places the RZZ complex in direct proximity to Mad1 residues 387-402 (Roscioli et al., 2020). This combined evidence strongly suggests that the Mad1^387Δ^ mutant is defective for an interaction with the RZZ complex. An interesting model for future study is that like Mad1’s interaction with Bub1, binding to the RZZ complex is facilitated by a core motif (387-402) but is multivalent in nature. Consistent with this, a recent report found that Cyclin B1 interacts with the most N-terminal region of Mad1 and facilitates its incorporation into the fibrous corona (Allan et al., 2020).

The RZZ pathway for Mad1 kinetochore recruitment has been studied for nearly a decade, yet the Mad1 region contributing to this behavior had not been identified previously.

This may have eluded researchers because Mad1 387-402 shows sequence conservation only in a small subset of eukaryotes. Multiple sequence alignments of putative Mad1 proteins from diverse eukaryotes (van Hooff et al., 2017), shows conservation of the Bub1 binding motif (RLK) but failed to identify homology near our region of interest (**Fig. S7A**). This is somewhat expected as the RZZ complex is not present in all eukaryotes, yet when alignments were limited to species with putative RZZ homologs conservation was still poor (**Fig. S7B**). In fact, strong conservation could only be observed among coelomate species like round worms, insects, and vertebrates (**Fig. S7C**). This suggests 387-402 is evolutionarily divergent relative to other Mad1 motifs and is further evidence that CRISPR tiling screens can identify regions not easily recognized by sequence homology. Consistent with this interpretation we find only weak conservation of Mad1 amino acids 387-402 among 100 vertebrates (Average PhyloP, **Fig. S5A**). Moreover, four of the essential uncharacterized domains we validated using cell biology (**Figs. 4, S5**) also show no statistical evidence for sequence conservation among vertebrates (**Fig. S5A**). Altogether these findings strongly suggest that our tiling screen is not limited by sequence homology and is an important tool for interrogating often ignored, evolutionarily divergent protein regions.

There are also limitations to the current tiling approach which should be considered. First, library coverage and domain resolution is partly determined by use of the “NGG” protospacer adjacent motif, required by type II CRISPR-Cas system from *Streptococcus pyogenes* (Mali et al., 2013). For example, large gaps in library coverage (148 nt maximum spacing) are due to a lack of NGG sequences (**Fig.1C**), which may score as false negatives. Utilizing CRISPR nucleases with a more permissive PAM sequence (e.g., xCas9 or Cas9-NG) (Hu et al., 2018; Nishimasu et al., 2018), should enhance tiling screens by allowing more uniform and closer spacing between sgRNAs. Second, in its current form, this approach will not identify regions for which a redundant gene exists (**Fig. 2C**). Third, while tiling mutagenesis appears robust when assaying aneuploid cell lines, gross genetic alterations (e.g., chromosome rearrangements, gene fusions, SNPs) may confound analysis of some genes (Munoz et al., 2016). Despite these limitations, we find that, in its current, form essential gene regions are readily identified; and there are a few if any false positives (e.g., “non-essential” regions identified as required for proliferation).

Altogether, this screening strategy is widely applicable and has benefits over other methodologies. Compared with traditional mutation screening, the cost and scale of tiling libraries are magnitudes more reasonable than chemical or UV induced mutagenesis strategies in human cells. Similarly, tiling mutagenesis targets endogenous genomic loci making it a better readout of cellular activity than libraries of mutant proteins expressed with highly active promoters from ectopic loci. Tiling mutagenesis screens are also an important advance beyond computational approaches that infer function based on sequence homology because tiling annotations are derived from phenotypic outcomes and thus ensure regions identified are truly important for protein function. Additionally, because sgRNA can be targeted nearly anywhere in this functional screen, important protein domains can be identified in regions resistant to homology-based analysis, namely disordered protein regions and rapidly evolving sequences.

## METHODS

### Mammalian cell culture

HeLa, ARPE^TERT^, ARPE^RAS^, HCT116, 293T, and HeLa FlpIn Cells (Etemad et al., 2015; Taylor et al., 1998) cells were grown in a high glucose DMEM (Thermo Fisher Scientific 11-965-118/Gibco 11965118) supplemented with antibiotic/antimycotic (Thermo Fisher Scientific 15240062) and 10% Fetal Bovine Serum (Thermo Fisher Scientific 26140095) at 37 °C supplemented with 5% CO2. For microscopy experiments, cells were seeded in 35mm wells containing acid washed 1.5 x 22mm square coverslips (Fisher Scientific 152222) and grown for 12-24 hours prior to transfections or immunostaining and most treatments are outlined in figures. Identity of each cell line was routinely validated by the presence of unique genetic modifications (Frt site, drug resistance genes, expression of transgenes) to ensure cross-contamination did not occur. Cell lines were also regularly screened for mycoplasma contamination using DAPI staining

To entirely depolymerize the microtubule cytoskeleton prior to immunofluorescence staining, cells were treated with 10 µM nocodazole (Sigma-Aldrich M1404) for one hour. To test SAC activity cells were instead treated with 500 nM nocodazole (Sigma-Aldrich M1404) for 20 hours prior to fixation.

### Library Design and cloning

All possible sgRNA target sequences within the protein coding DNA sequence (CDS) of 48 target genes were identified using the Broad Institutes GPP sgRNA Designer (since redesigned as CRISPick) by inputting gene symbols (Doench et al., 2016; Sanson et al., 2018). CRISPick output sequences for every unique spCas9 PAM (NGG) which cut within the CDS of the consensus or longest transcript for each gene. An initial library was generated from sgRNA that uniquely targeted the gene of interest and no other exonic regions within the genome. To prevent bias in the library near poly-G repeats, sgRNA were removed to ensure a minimum spacing of 5 nt between cut sites within the CDS. In regions where there were >50 nts between sgRNA, we included sequences with limited off-target sites to increase the resolution of screening. One exception being last minute additions, CENP-S and CENP-X where all unique sgRNA were included with no adjustment for minimum or maximum spacing between sgRNAs. A pooled single stranded DNA 60-mer library containing all sgRNA sequences was synthesize by Twist Biosciences. Oligomers were designed with a universal 20 nucleotides flanking the 5’ and 3’ with unique sgRNA sequences in the middle 20 nucleotides. The library was PCR amplified using universal primers that annealed to the common flanking sequence and appended homologous sequences at 5’ and 3’ ends of the PCR product to enable Gibson assembly (New England Biolabs E2611) into pZLCv2_puro_1KF. The vector pZLCv2_puro_1KF was linearized by digestion with restriction enzyme Esp3I and both PCR product and vector were gel purified prior to assembly.

### CRISPR/Cas9 Screening

Outgrowth screens were performed as previously described (Toledo et al., 2015). The library of sgRNA-containing donor plasmids, pPAX2, and pMD2.G were co-transfected into 293T cells using polyethyleneimine (PEI, Polysciences 23966-1). Virus-containing supernatant media were harvested 48 hours post transfection and passed through 0.45 µm filters, concentrated by centrifugation, and stored at -80 °C. Each cell line was infected with varying volumes of concentrated virus in the presence of polybrene (Sigma Aldrich 107689) to determine the concentration that conferred survival in puromycin to 30% of cells, representing an MOI of 0.3 where a single infection per cell is the most likely outcome. Three replicates of each cell line were infected at scale to ensure 650x representation of the library and then 24 hours later were exposed to 1 µg/mL puromycin. 72 hours after infection the puromycin containing media was replaced with drug free media. 96 hours after infection cells were trypsinized and re-seeded to maintain 650x representation, while excess cells were harvested as an initial timepoint. Over the next eight days replicates were sub-cultured to maintain representation and eventually harvest a final population. Genomic DNA was extracted from 5 million cells (∼650x representation) in the initial and final populations each using a QiaAMP DNA Blood Purification Mini Kit (Qiagen 51104) and then sgRNA sequences were amplified from each sample using a two-step PCR. For the first step, a 12 cycle PCR was performed using Phusion polymerase (New England Biolabs M0530) to amplify from all the genomic DNA extracted from the 5 million cells per sample (70- 80 reactions). For the second step, an 18 cycle PCR was amplified from the pooled first step using primers coding 6 bp Illumina sequencing barcodes used for multiplexing biological samples. The final amplicon was purified from genomic DNA using a Monarch PCR and DNA Cleanup Kit (New England Biolabs T1030) and quantified with a Qubit 2.0 Fluorometer. Samples were then sequenced using an Illumina HiSeq 2500. Deconvoluted sequencing results have been submitted to NCBI’s GEO repository under the submission record GSE179188 (https://www.ncbi.nlm.nih.gov/geo/query/acc.cgi?acc=GSE179188) and can be accessed by reviewers using the token ‘qfolmasydxahjyb’, which will be made publicly available at the time of publication.

### Computational Analysis of Tiling Data

Relative changes to the amount of sgRNA sequence detected in final versus initial samples were determined by the CRISPR-SURF package run from the command line (https://github.com/pinellolab/CRISPR-SURF) (Hsu et al., 2018). The SURF package includes ‘CRISPR-SURF Count’ which outputs logFC values for each sgRNA within the library. This output was used by CRISPR-SURF to deconvolve tiling data and identify the targeted genomic regions that had a negative effect on proliferation relative to non-targeting controls. This output was also the input for ProTiler (https://github.com/MDhewei/protiler) (He et al., 2019) and TiVex and was used to calculate Z scores. In a few instances data were excluded from computational analysis. CRISPR-SURF Count did not report values for sgRNA containing a TTTT repeat due to their likelihood of causing premature transcriptional termination. No other data were removed from global lists but in the case of genes BIRC5 and KNL1 we generated the library using transcripts containing rare or mutually exclusive exons and when analyzing them at the protein level we mapped results to a more common transcript that does not contain those regions.

<u>Ti</u>ling data with a con<u>vex</u> fused lasso (TiVex) analysis built upon previous approaches for analyzing tiling data that used a Fused Lasso to deconvolve complex signals. The Fused Lasso optimizes the cost function 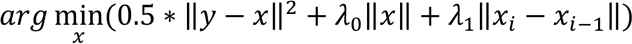 (Hsu et al., 2018; Tibshirani and Taylor, 2011), but this was designed for sparse regulatory elements, while functional motifs in proteins are large blocks and may cover a large portion of proteins. If the sparsity induced penalty is reduced (1_#_ = 0), then the cost function is equivalent to identifying segmentations and not useful. To balance sparseness we used a Convex Fused Lasso (Parekh and Selesnick, 2015) to deconvolve the data. This approach optimizes the cost function, 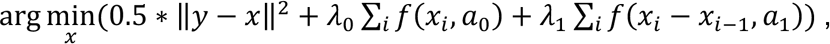 where 8(. ) is a transform function and 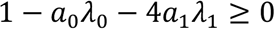 defining the convex shape in the transformed space. TiVex regions were identified as negatively enriched by comparing the per gene signal to a global average of all genes. Code for TiVex analysis will be made publicly available upon publication at https://labs.icahn.mssm.edu/zhulab/software/ or https://github.com/integrativenetworkbiology

We used CRISPR SURF and TiVex to identify protein domains in our datasets consisting of 4 different cell lines. The domains identified were called “Positive Domains”. To calculate “Negative Domains“, we divided up the regions which were not identified as a Positive Domain by CRISPR SURF or TiVex into 3 or 45 AA windows respectively for the 48 genes that were tiled. Next, we simulated datasets with low efficiency sgRNAs by randomly transforming x% (10, 20, 30, 40, and 50%) of sgRNAs targeting a gene from their true value to a NTC value. This was performed 10 times at each % resulting in 50 datasets. For each of these datasets, we used CRISPR-SURF or TiVex to identify protein domains. Domains of min width of 3 AA and FDR < 0.05 were considered in our analysis. For each percentage of signal lost, we pooled results across 10 runs and calculated True positive (TP), False Positive (FP), True Negative (TN) , False Negative (FN), Precision, Recall and F1 score. True Positives (TP) is the number of domains retrieved from simulated data that overlapped with a “Positive domain”. True Negative (TN) is the number of negative domains which were correctly classified as a negative domain in the simulated data and were also a “Negative domain”. False Positive (FP) is the number of domains that were identified when low efficiency sgRNAs were simulated but were a “Negative domain” in the original dataset. False Negative represents the positive domains which were not identified in the simulation. Precision is the fraction of correct domains among the total predicted domains calculated as: Precision ={TP}/{TP+FP}. Recall is the fraction of correct domains among the total domains in the dataset. It indicates how many total domains of the actual dataset were picked up as a domain with noise and is calculated as Recall ={TP}/{TP+FN}. F1 score integrates Precision and recall and is calculated as F1 score = 2* ((precision*recall)/(precision+recall)).

### Statistics

Outside of tiling analysis packages, GraphPad Prism version 9.1.0 was used for statistical analysis. Each test (paired, multiple comparison, etc.) is specifically identified in figure legends, and generally error bars represent 95% confidence intervals.

### Generation of Modified Human Cell Lines

HeLa FlpIn Trex cells encoding wild type and mutant proteins were generated as previously described (Herman et al., 2020). Briefly, HeLa FlpIn Trex cells were transfected with Flp recombinase (p0G44) and a donor plasmid encoding the protein of interest using lipofectamine 2000 (Invitrogen 11668027) according to manufacturer instructions or PEI (Polysciences 23966-1). 48 hours post transfection, media was supplemented with 500 µg/mL hygromycin (Invitrogen 10687010) and cells were negatively selected for 3 days. Expression of EGFP fusion proteins was then induced by addition of 1 µg/mL doxycycline (Sigma-Aldrich, D9891) and EGFP expressing cells were positively selected by FACS. Doubly selected polyclonal populations were frozen and stored for future experiments.

ARPE^TERT^ and ARPE^RAS^ were generated through single or serial retroviral transductions followed by selecting for cells expressing a single drug resistance gene as previously described (Kendall et al., 2005; Toledo et al., 2014) using methods like the lentiviral particle generation detailed above for the CRISPR/Cas9 tiling library.

### Nucleic Acid Reagents

Mad1 and Sgo1 FRT/TO/Hygro vectors were a gift from Jennifer DeLuca. The coding DNA sequence for Mad1 and Sgo1 was amplified from cDNA libraries and thus for proliferation retests synthetic sgRNA targeting these genes span intron-exon boundaries to ensure the ectopic copy was not targeted. All other coding sequences were generated as codon optimized and thus sgRNA resistant gBlocks (IDT) and inserted into restriction enzyme linearized pcDNA5 FRT/TO/Hygro by Gibson assembly and sequence verified.

### sgRNA:Cas9 mediated gene knockout

Genes were knocked out using 1-2 synthetic sgRNAs (Synthego) in complex with spCas9 (Aldeveron 9214) that were electroporated into cells using a nucleofector system (Lonza V4XC- 1032) according to published methods (Hoellerbauer et al., 2020a; Hoellerbauer et al., 2020b). Briefly, 120k cells were mixed with either targeting or non-targeting sgRNA:Cas9 complexes in complete SE nucleofector solution. Cell solutions were added to 16-well mini-cuvettes cells and electroporated using program CN-114. Cells were split into two well, one with doxycycline and one without, then cell numbers were assayed 5-7 days later.

### Immunopurification

T-225 flasks of Mad1^WT^, Mad1^Δ387^, Mad1^R617A^, and Mad1^Δ387+R617A^ HeLa FlpIn Trex cells were grown to 50% confluence and induced with 1 µg/mL of doxycycline for 16-20 hours. Cells were then treated with 5 µM nocodazole and 10 µM MG132 for four hours to elicit a robust mitotic arrest with unattached kinetochores. Cells were harvested by trypsin digestion, counted by hemocytometer, and then centrifuged. The cell pellet was resuspended in 1 µL of complete lysis buffer (25 mM HEPES, 2 mM MgCl2, 0.1 mM EDTA, 0.5 mM EGTA, 15% Glycerol, 0.1% NP-40, 150 mM KCl, 1 mM PMSF, 1 mM sodium pyrophosphate, 1x Pierce Protease Inhibitor Cocktail [Thermo Scientific 88666]) for each 200,000 cells, and then snap frozen in liquid nitrogen. Samples were thawed and sonicated with a CL- 18 microtip for 15 s at 50% maximum power with no pulsing two times using a Fisher Scientific FB50 sonicator. Approximately 150 U of Benzonase nuclease (Millipore E1014) was added to samples and incubated at room temperature for 15 min. The samples were centrifuged at 16,100 x g at 4°C in a tabletop centrifuge for 30 minutes. Clarified lysates were moved to fresh microfuge tubes and 60 mL of Protein G Dynabeads (Thermo Fisher Scientific 10009D) conjugated with anti-EGFP monoclonal antibody (Sigma Aldrich 11814460001) or anti-Zw10 polyclonal antibody (Proteintech 24561-1-AP) as previously described (Akiyoshi et al., 2009) were added to 250 µL of lysate and incubated at 4°C with rotation for 90 min. Beads were washed six times with lysis buffer lacking PMSF, sodium pyrophosphate, and protease inhibitor cocktail. Proteins were eluted from beads in 60 mL of 1x SDS sample buffer and incubated at 95°C for 10 minutes.

### Immunoblotting

Expression of Flag- and EGFP-tagged proteins was induced with media containing 1 µg/mL doxycycline (Sigma-Aldrich D9891) 12-24 hours prior to harvesting. Cells were isolated via trypsinization and then centrifuged. Immunoblotting was performed as previously described (Herman 2020). Cells previously exposed to doxycycline to induce wild type or mutant protein expression were harvested by trypsinization then resuspended in complete lysis buffer and frozen in liquid nitrogen. Samples were thawed and sonicated with a CL-18 microtip for 20 seconds at 50% maximum power with no pulsing three times using a Fisher Scientific FB50 sonicator. Benzonase nuclease (Millipore E1014) was added to samples and incubated at room temperature for five minutes, then samples were centrifuged at 16,100 x g at 4 °C in a tabletop centrifuge. Relative protein concentrations were determined for clarified lysates, and samples were normalized through dilution. Denatured samples were run on Tris buffered 10 or 12% polyacrylamide gels in a standard Tris-Glycine buffer. Proteins were transferred to a 0.45 µm nitrocellulose membrane (BioRad 1620115) for two hours at 4 °C in a transfer buffer containing 20% methanol. Membranes were washed in PBS+0.05% Tween-20 (PBS-T) and blocked with PBS-T+5% non-fat milk and incubated with primary antibodies overnight at 4 °C. Antibodies were diluted in PBS-T by the following factors or to the following concentrations: anti-GAPDH clone 6C5 (Millipore Sigma MAB374) 1 µg/mL; anti-GFP clone JL-8 (Takara 632381) 0.5 µg/mL; anti-Flag clone M2 (Sigma Aldrich F3165) 2 µg/mL, anti. HRP conjugated anti-mouse secondary antibodies (GE Lifesciences NA931) were diluted 1:10,000 in PBS-T and incubated on membranes for 45 minutes at room temperature. Immunoblots were developed with enhanced chemiluminescence HRP substrates SuperSignal West Dura (Thermo Scientific, 34076) using a ChemiDoc MP system (BioRad).

### Immunofluorescent Staining

Upon completion of experimental manipulations, cells grown on coverslips were immediately chemically crosslinked for 15 minutes with 4% PFA diluted from a 16% stock solution (Electron Microscopy Sciences, 15710) with 1x PHEM (60 mM PIPES, 25 mM HEPES, 5 mM EGTA, 8 mM MgSO4) or 1x PHEM+0.5% TritonX100. Coverslips were washed with 1x PHEM+0.5% TritonX100 for 5 minutes, then washed 3 more times with 1x PHEM + 0.1% TritonX100 over 10 minutes. Cells were blocked for 1-2 hours at room temperature in 20% goat serum in 1x PHEM. Anti-centromere protein antibody or ACA (Antibodies Inc. 15-235) was diluted in 20% goat serum at a 1:600 dilution factor. Coverslips were incubated overnight at 4°C in the primary antibody, then washed four times with 1x PHEM + 0.1% TritonX100 over 10 minutes. Goat anti-human secondary antibodies conjugated to AlexaFluor 647 (Invitrogen) were diluted at 1:300 in 20% boiled goat serum. Coverslips were washed four times with 1x PHEM + 0.1% TritonX100 over 10 minutes, then stained for one minute with 30 ng/mL 4′,6-diamidino-2-phenylindole (DAPI, Invitrogen D1306) in 1x PHEM. Coverslips were washed two times with 1x PHEM, then immersed in mounting media (90% glycerol, 20 mM Tris [pH= 8.0], 0.5% w/v N-propyl gallate) on microscope slides and sealed with nail polish.

### Microscopy and Image Analysis

Fixed cell images were acquired on either a Deltavision Elite or Deltavision Ultra deconvolution high-resolution microscope, both equipped with a 60x/1.42 PlanApo N oil-immersion objective (Olympus). Slides imaged on the Elite were collected with a Photometrics HQ2 CCD 12-bit camera, while those imaged on the Ultra were equipped with a 16-bit sCMOS detector. On both microscopes, cells were imaged in Z-stacks through the entire cell using 0.2 µm steps. All images were deconvolved using standard settings. Z projections of the maximum signal in the ACA or EGFP channel were exported as TIFFs for analysis by Cell Profiler 4.0.7 (29969450). ACA images were used to identify regions of interest after using a global threshold to remove background signal and distinguishing clumped objects using signal intensity. The signal intensity within these regions was quantified from the EGFP images, and then for background correction the regions were expanded by one pixel along the circumference and signal intensity was again quantified on the EGFP channel. Background intensity was found by subtracting the intensity of the original region from the 1-pixel expanded region. The background intensity per pixel was quantified by dividing the background intensity by the difference in area between two regions. This was then multiplied by area of the original object and subtracted from the intensity of the original object. The mean value per image was then determined and displayed in figures. Representative images displayed from these experiments are projections of the maximum pixel intensity across all Z images. Photoshop was used to crop, make equivalent, linear adjustments to brightness and contrast, and overlay images from different channels.

### Computational Analysis of Protein Sequences

#### Databases of Conserved and Disordered Proteins

PhyloP scores for nucleotides within the genomic regions corresponding to essential regions identified by tiling were downloaded manually from UCSC genome browser and mean values calculated. Disordered regions were identified using the D2P2 database (Oates et al., 2013) based on the agreement of >75% of disorder prediction algorithms. Our comprehensive list of Pfam domains were identified using gene IDs.

#### Multiple Sequence Alignments

Eukaryotic Mad1 orthologs were identified previously (van Hooff et al., 2017) and a multiple sequence alignment of the entire proteins was generated with ClustalOmega (Sievers et al., 2011) default parameters and displayed in JalView 1.8 (Waterhouse et al., 2009).

## ACKNOWLEDGEMENTS

We thank members of the Biggins and Paddison Labs for helpful discussions, and Bruce Clurman for providing HCT116 cells and Jennifer DeLuca for plasmids encoding SGO1 and MAD1L1. This work was supported by the: American Cancer Society (ACS-RSG-14-056-01) (PP); NIH (R01CA190957; R01NS119650, P30CA15704) (PP); NIH (R01GM064386) (SB); and Robert J. Kleberg, Jr. and Helen C. Kleberg Foundation (PP). SB is an investigator of the Howard Hughes Medical Institute.

## AUTHORS’ CONTRIBUTIONS

Conceptualization, JH, SB, and PP; Methodology, JH, JZ, SB, PP; Validation, JH, LC; Investigation, JH ; Formal Analysis, JH, SA, JZ, and PP; Writing - Original Draft, JH, JZ, SB, and PP; Writing - Review & Editing, JH, SB, and PP; Visualization, JH, SA, JZ, PP; Supervision, JH, SB, and PP; Funding Acquisition, SB and PP.

---

**Figure S1.**
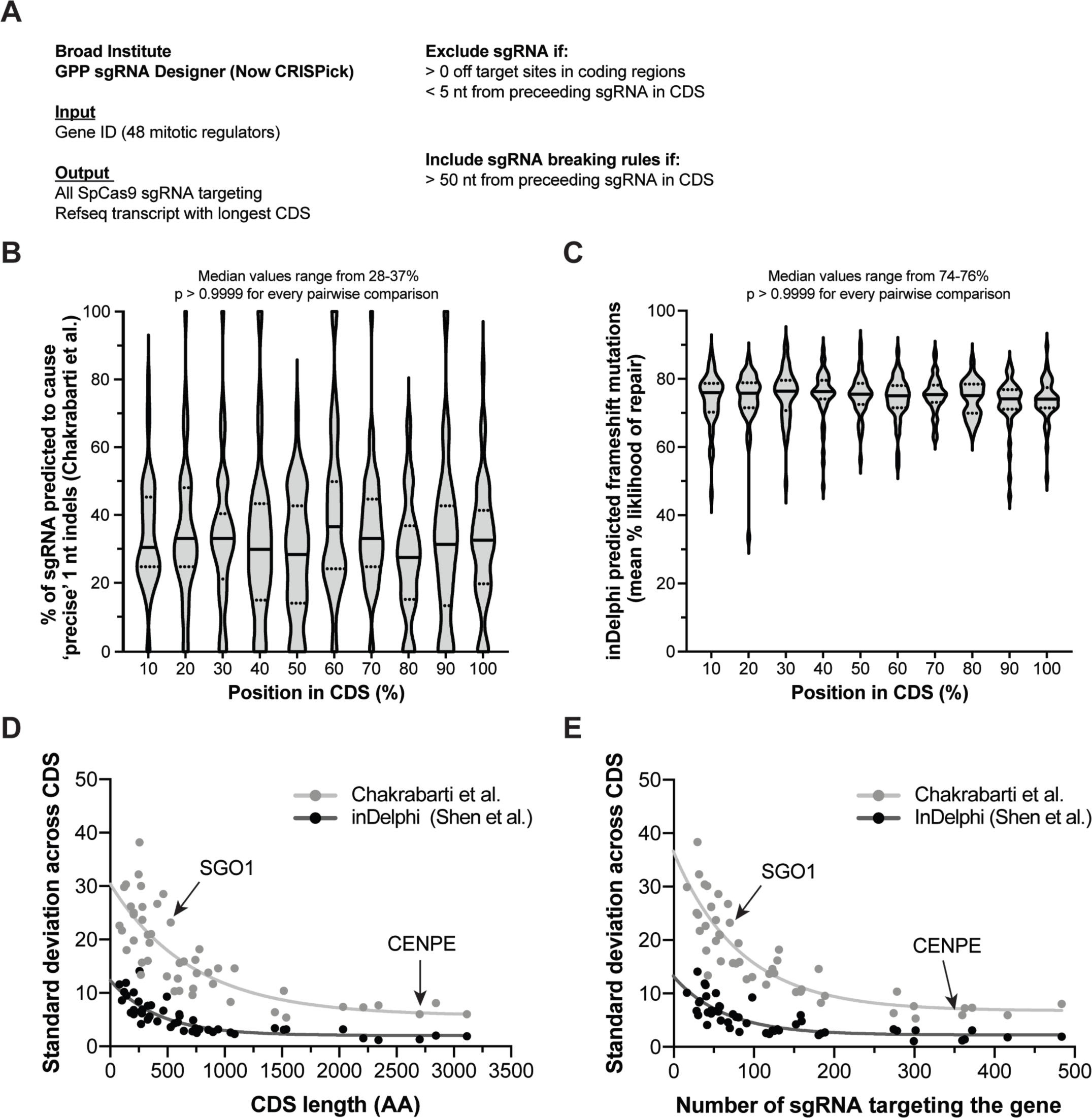
Tiling library design strategy benefits from a lack of positional repair bias. (A) Scheme for choosing sgRNA included in the library. (B-C) All sgRNAs in the library were binned based on the position they target (% of CDS). (B) For each bin, in each gene, the percent of sgRNA predicted to cause 1nt indels based on sequence constraints (Chakrabarti et al., 2019) are plotted. Violin plots show median (solid lines), quartiles (dotted lines) and range of bins for all genes. (C) For each bin, in each gene, the mean percent likelihood of forming a frameshift mutation based on sequence constraints (Shen et al., 2018) are plotted. Violin plots show median (solid lines), quartiles (dotted lines) and range. (D-E) The standard deviation across each bin in (B) and (C) for each gene was calculated and is plotted as a single dot. Deviations are plotted relative to the CDS length (D) or the total number of sgRNAs targeting the gene (E). Data are fitted with a single-phase exponential decay curve.

**Figure S2.**
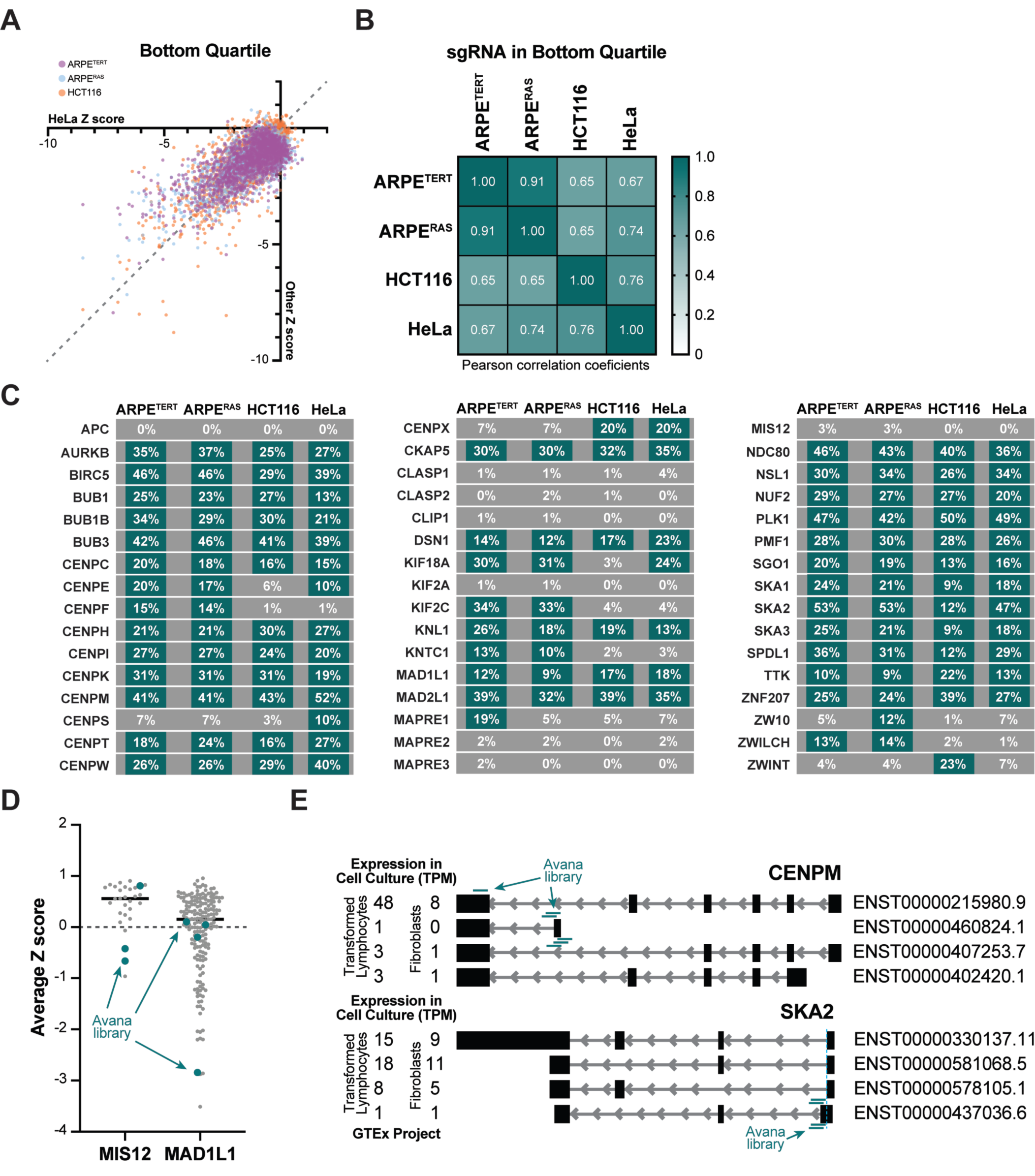
Tiling library is reproducible and differences from genome-wide data have biological causes. (A) Each sgRNA’s average Z score from three replicates of ARPE^TERT^, ARPE^RAS^, and HCT116 cells are plotted relative to each sgRNA’s average Z score from three replicates of HeLa cells. These data exclude non-targeting controls and only show sgRNA in the bottom quartile of Z-scores from HeLa cells. The dashed line (y = x) is plotted for reference. (B) Correlation matrix and heatmap for the bottom quartile of targeting sgRNA in each cell line with Pearson correlation coefficients displayed. (C) Percent of sgRNA with Z < -1.0 targeting each gene are shown for all four cell lines. Cell lines meeting the >8% cutoff are highlighted in teal. (D) Z scores for each sgRNA targeting genes where disagreement between tiling and DepMap data disagree (MIS12 or MAD1L1), with sgRNA sequences present in both libraries colored teal. (E) Numerous transcript models of CENPM and SKA2 (right) along with their relative expression in cultured cells from public RNA-seq datasets (right). The location targeted by each sgRNA in the DepMap (Avana) library shown as teal bars indicating they target rarely used exons.

**Figure S3.**
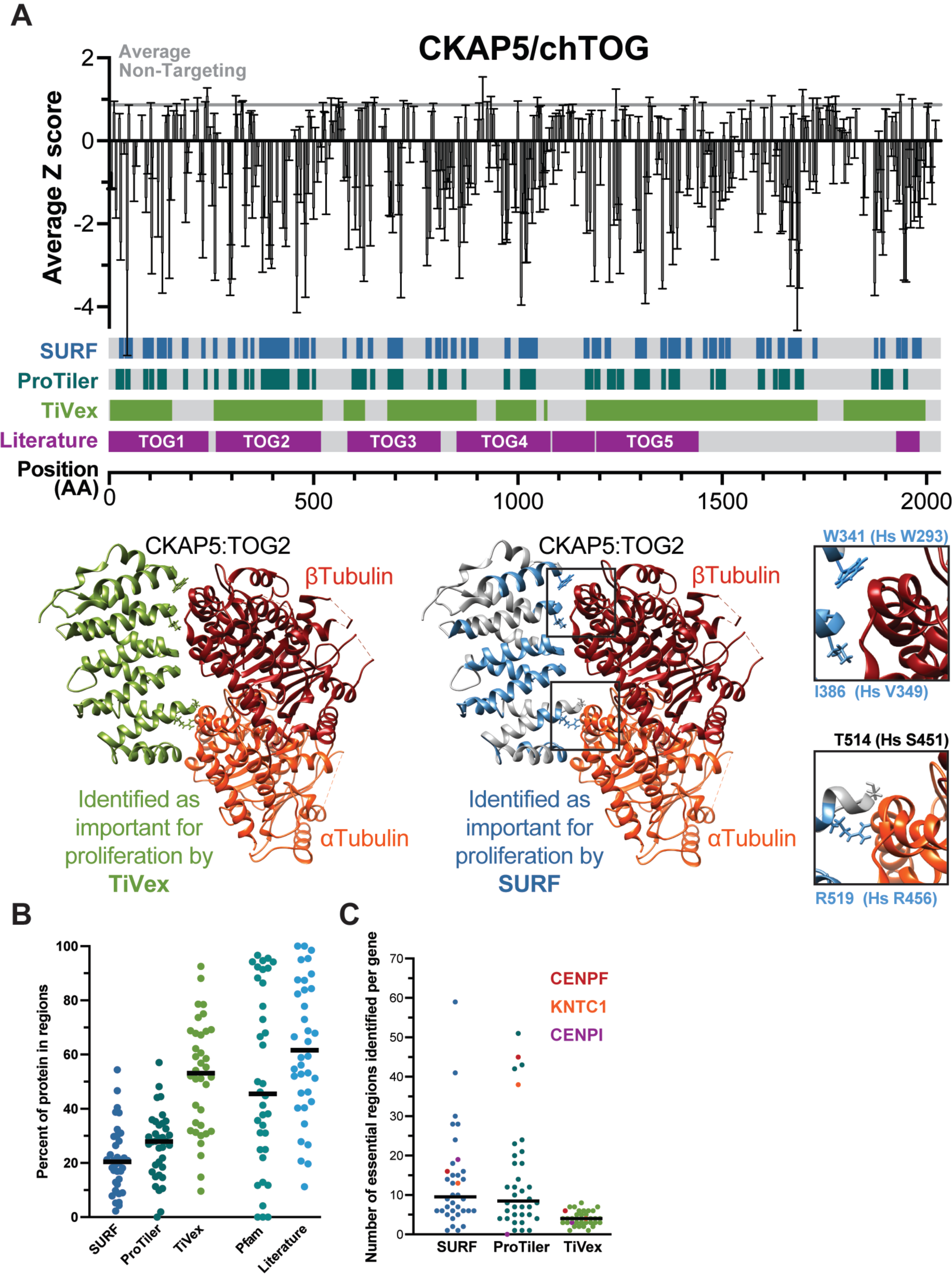
More evidence of SURF, ProTiler, and TiVex resolution. (A) (top) Tiling profile for CKAP5/chTOG as shown in Fig. 3. (bottom) Crystal structure of TOG2 (∼aa250-500) from budding yeast homolog bound to a tubulin dimer (PDB: 4U3J) are colored based on the homologous regions identified by TiVex (green) and SURF (blue) as important for proliferation (Ayaz et al., 2014). Four residues absolutely required for the protein-protein interaction are shown in blowups. (B) Percent of each protein sequence identified by the three analysis methods or Pfam or within literature. Average values reported in Fig. 3B. (C) The number of regions identified per gene by each analysis method. Specific genes showing key differences between SURF and ProTiler are colored uniquely in all three columns.

**Figure S4.**
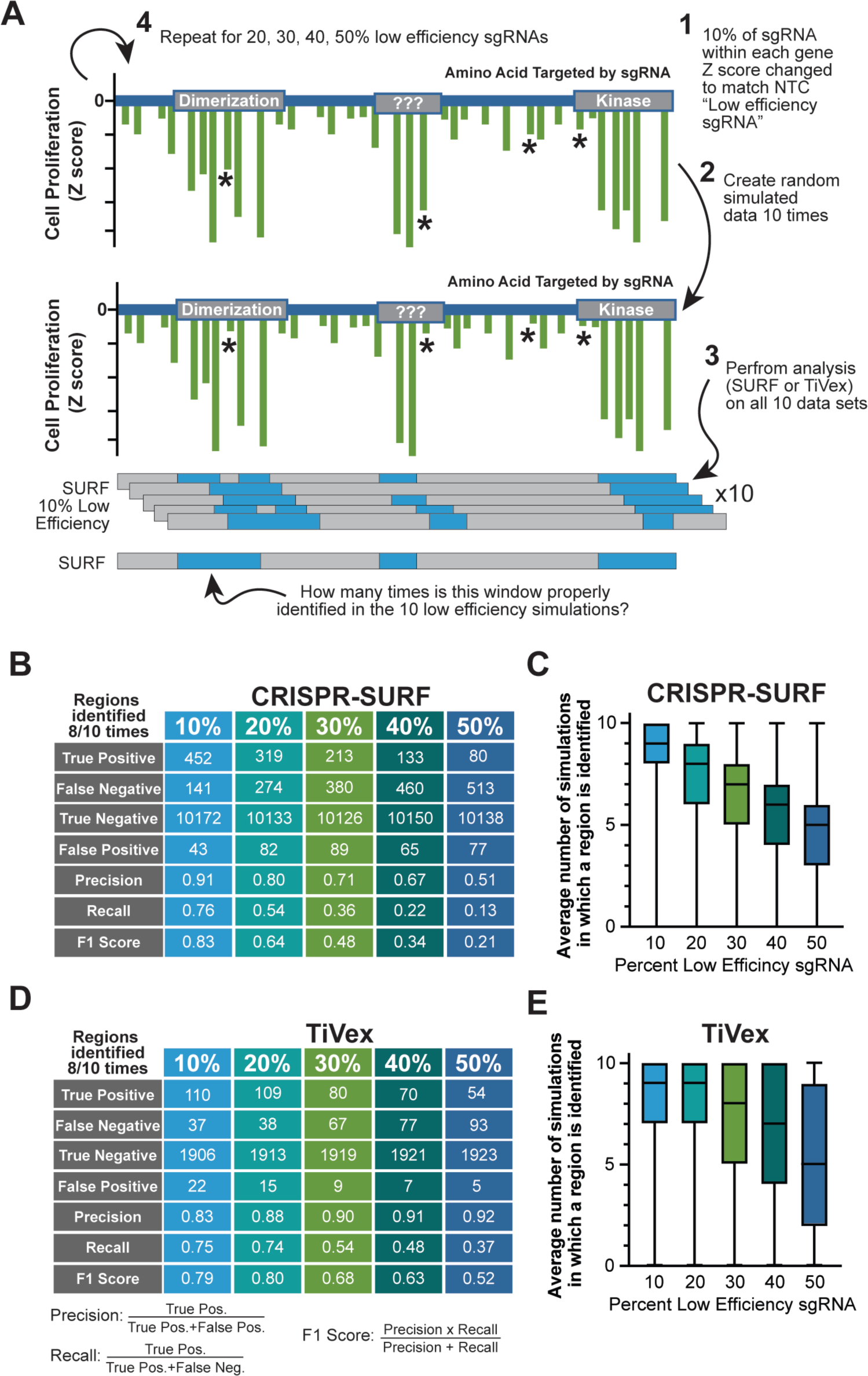
Simulation to determine how low efficiency sgRNA affect tiling analysis. (A) Schematic showing how datasets simulating 10, 20, 30, 40, or 50% low efficiency sgRNAs were generated for all 48 mitotic factors, which were analyzed 10 times each using CRISPR-SURF or TiVex. (B) Table representing the ability of simulated data to recapitulate original findings of CRISPR SURF analysis among all kinetochore genes, where ‘positives’ are identified in 8/10 simulations. (C) Box and whisker plot showing how often individual SURF regions were identified datasets simulating low efficiency sgRNAs. (D,E) Same analysis as B and C but for TiVex. Boxes in C and E represent median and quartiles while whiskers show range of datapoints.

**Figure S5.**
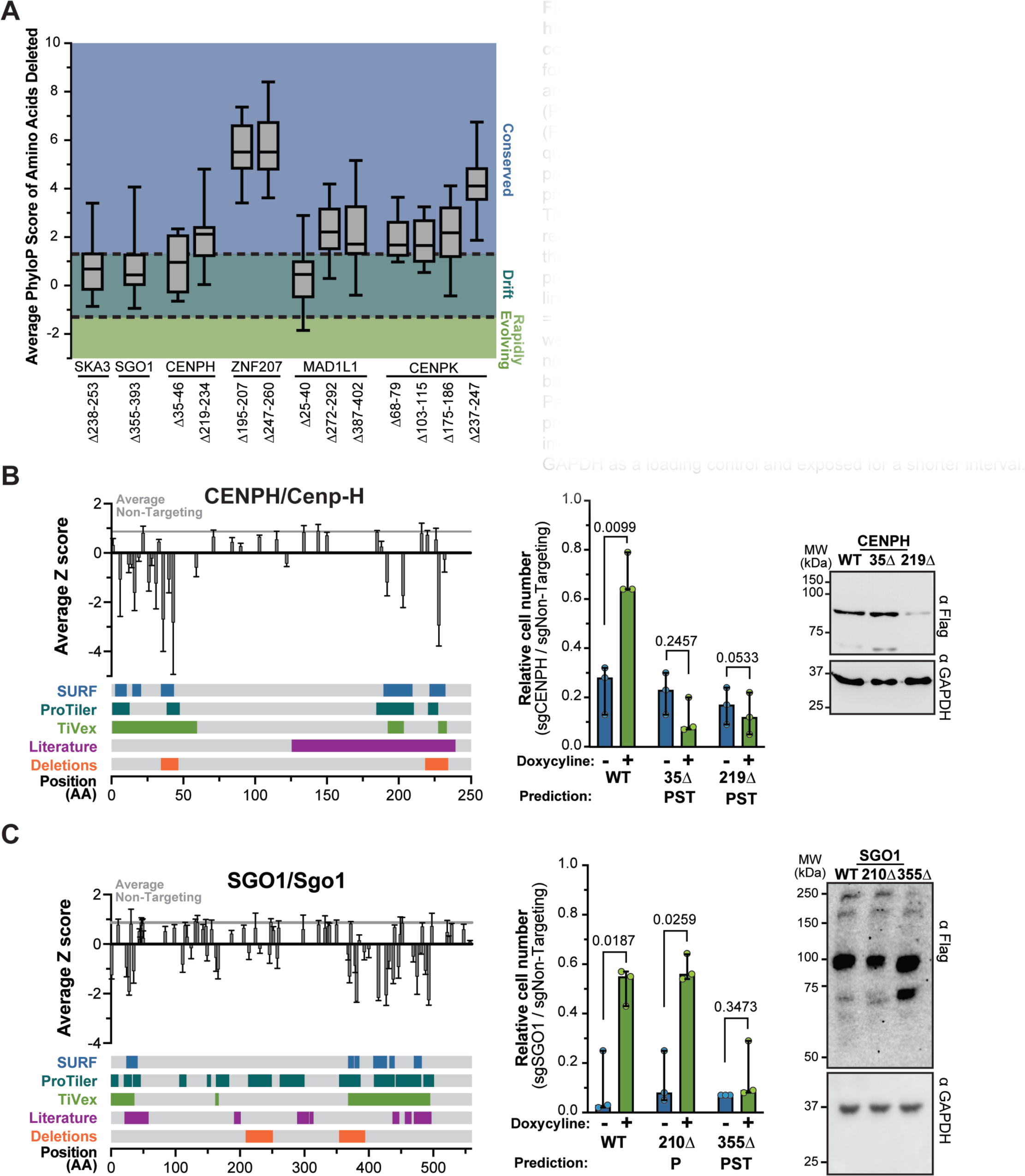
Functional validation and characterization of 11 high-resolution regions within 5 genes identified by tiling, continued. (A) Average PhyloP scores across 100 vertebrates for each codon within deletion mutants from figures 4 and S5 are plotted. Dashed lines denote sequences statistically (P<0.05) unchanged (PhyloP > 1.3) or rapidly changing (PhyloP < -1.3) sequences. Boxes represent median and quartiles while whiskers show range of datapoints. (B-C) Tiling profile, validation of proliferation phenotype, and assay of protein stability for (B) CENPH/Cenp-H and (C) SGO1/Sgo1. Tiling profiles are the same as Fig. 3 while also showing regions that were deleted. Cell proliferation was assayed as the cell number after knockout of endogenous protein in the presence (green) or absence (blue) of doxycycline in each cell line. The overlap of deletion and analysis methods shown as P = ProTiler, S = CRISPR-SURF, and T = TiVex. Cell numbers were normalized to the same cell line electroporated with a non-targeting control. Each dot is a biological replicate with bars showing median values and 95% confidence intervals. Paired t tests were used to determine P-values. Steady state protein levels of wild type and mutant proteins was assayed by immunoblot in the presence of endogenous protein, using GAPDH as a loading control and exposed for a shorter interval.

**Figure S6.**
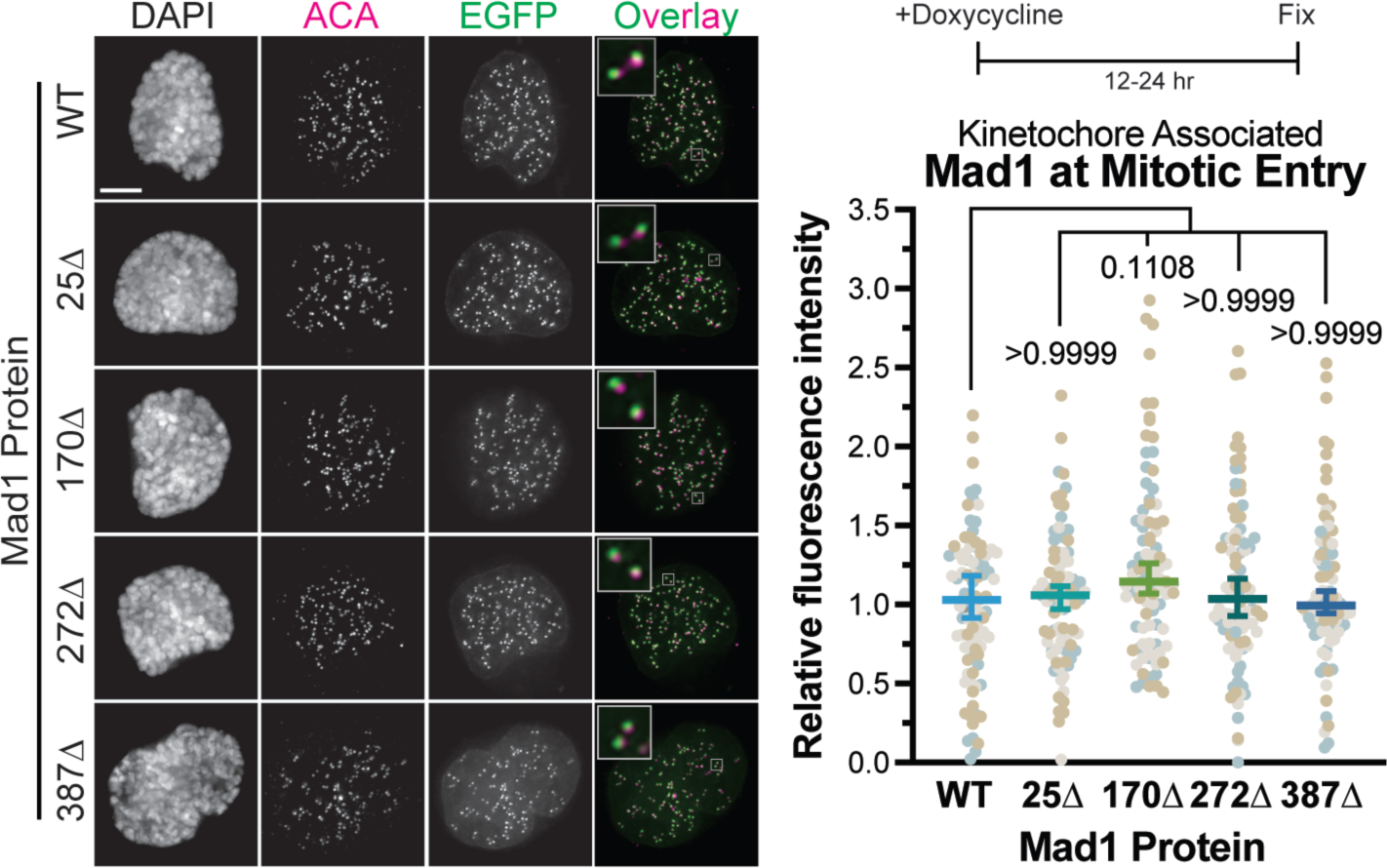
Mutations in Mad1 do not affect localization to kinetochores immediately following nuclear envelope breakdown. Kinetochore association of EGFP-Mad1 wild type and mutant fusion proteins was determined by the EGFP proximal to anti-centromere antibody (ACA) in the presence of endogenous Mad1 immediately after nuclear envelope breakdown. Representative images on left with quantifications on right. Each dot represents the average kinetochore signal from a single cell, cells from three biological replicates are colored differently. Dunn’s multiple comparisons test was used to determine P-values. Scale bars are 1 µm, all averages and error bars in figure are median values and 95% confidence intervals.

**Figure S7.**
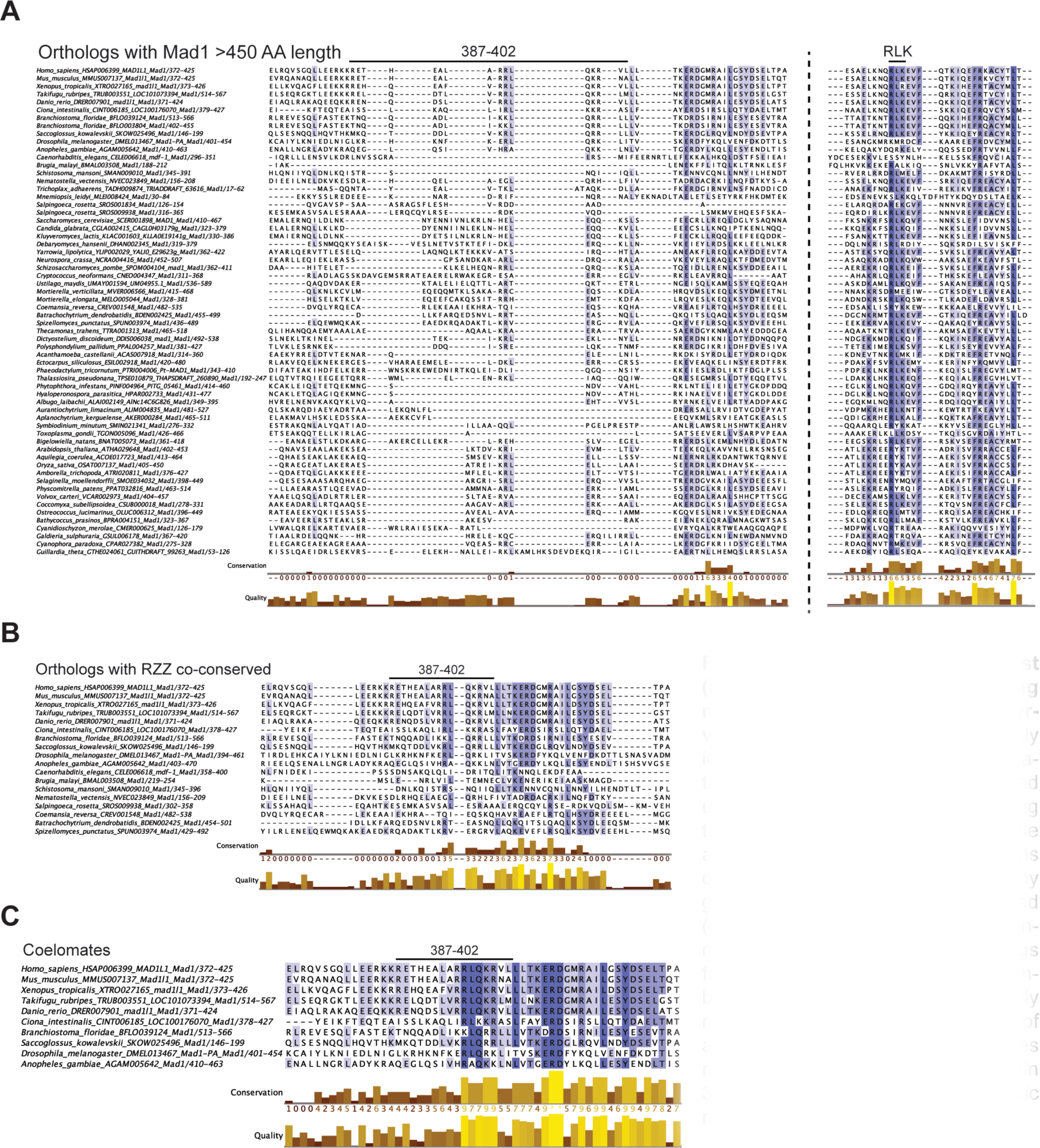
Mad1 region of interest (aa 387-402) is divergent among most eukaryotes but shows conservation in coelomates. (A) Previously identified (van Hooff et al., 2017) putative Mad1 orthologs were aligned using ClustalO and colored according to percent sequence identity. Multiple alignment was driven by other regions of the protein and 387-402 had many gaps (left), while RLK motif is identified (right). (B) Multiple sequence alignment was restricted to Mad1 orthologs from species that also encode members of the RZZ complex (primarily metazoan). (C) Further restriction of alignment to only coelomate species revealed a conserved sequence within 387-402 that is enriched for basic residues.

## Description of Supplementary Tables

**Supplementary Table 1**: sgRNA sequences, targets, raw, and normalized counts from Illumina sequencing.

**Supplementary Table 2**: Tiling regions identified using CRISPR SURF, ProTiler, and TiVex in 48 kinetochore genes

**Supplementary Table 3**: Annotations within 48 kinetochore genes from D2P2 and Pfam databases and manually curated from published literature.

**Supplementary Table 4**: Key resources used for this study.

